# DNA-targeting short Argonaute triggers effector nuclease to protect bacteria from invaders

**DOI:** 10.1101/2023.06.08.544223

**Authors:** Maria Prostova, Anna Kanevskaya, Vladimir Panteleev, Lidia Lisitskaya, Kristina V. Tugaeva, Nikolai N. Sluchanko, Daria Esyunina, Andrey Kulbachinskiy

## Abstract

Two prokaryotic defence systems, Argonautes (pAgos) and CRISPR-Cas, detect invader nucleic acids using complementary guides. Upon recognition, the target is cleaved through nuclease activities of pAgo or Cas proteins thus protecting the cell from invasion. However, not all pAgos are active nucleases. Members of a large clade of short pAgos bind nucleic acid guides but lack nuclease activity suggesting a different mechanism of action. Here, we have investigated short pAgo from *Novosphingopyxis baekryungensis* (NbaAgo). We have shown that NbaAgo forms a heterodimeric complex, SPARDA, with a co-encoded effector nuclease. RNA-guided target DNA recognition unleashes the nuclease activity of SPARDA leading to indiscriminate collateral cleavage of DNA and RNA. Activation of SPARDA results in cell death during plasmid transformation or phage infection, thus protecting bacterial population from invaders. The collateral activity of SPARDA allows highly sensitive detection of specific DNA targets. SPARDA expands the list of prokaryotic immune systems that elicit suicidal cell response with a unique range of nuclease activities, creating additional opportunities for biotechnologies.

## Introduction

Argonautes are a ubiquitous family of proteins that can be programmed for recognition of specific nucleic acid targets with short guide oligonucleotides. Argonautes were first discovered in eukaryotes due to their central role in RNA interference, but subsequent analysis demonstrated that they originated in prokaryotes ^1-8^. Unlike eukaryotic Argonautes (eAgos), most studied prokaryotic Argonautes (‘long’ pAgos, containing the N, L1, PAZ, L2, MID and PIWI domains) were shown to recognize and cleave DNA targets, using short guide DNAs or RNAs^9-16^. In contrast, ‘short’ pAgos, which form the largest group of pAgos, contain only two conserved domains, MID and PIWI, and lack catalytic residues found in the PIWI domain in long pAgo nucleases ^3-5,7,17^. Recent studies showed that short pAgos are usually co-encoded with proteins containing an APAZ domain (for <u>a</u>nalog of <u>PAZ</u>, not homologous to PAZ) fused to an additional effector ^3,5,17,18^. Studied short pAgos form binary complexes with their partner proteins, in which APAZ likely completes the pAgo structure to form the nucleic acid binding cleft ^17-20^. Recognition of target DNA by short pAgos can activate the associated effector domain, SIR2 or TIR, leading to hydrolysis of cellular NAD^+^ and cell death, thus preventing plasmid transformation or viral infection ^18,20,21^. However, the vast majority of short pAgo systems remain unstudied. The mechanisms of discrimination of invader DNA, the diversity of effector domains and their activation pathways are unknown for most short pAgos.

## Results

### SPARDA systems encode a predicted effector nuclease

To investigate the diversity of effectors associated with short pAgo systems, we performed phylogenetic analysis of APAZ domains and co-encoded effector domains in prokaryotic genomes. APAZ domains form four major clades, S1A, S1B, S2A and S2B (Fig. 1A) ^5,17,18^. Clades S1A and S1B correspond to SPARSA systems (for <u>s</u>hort <u>p</u>rokaryotic <u>Ar</u>gonaute and <u>S</u>IR2-<u>A</u>PAZ), consisting of short pAgos and associated SIR2-APAZ proteins. Clade S2A includes TIR-APAZ proteins (or Mrr-TIR-APAZ) from SPARTA systems (for <u>s</u>hort <u>p</u>rokaryotic <u>Ar</u>gonaute and <u>T</u>IR-<u>A</u>PAZ). Clade S2B is most diverse and includes a subgroup of SIR2 effectors, a subgroup of TIR effectors, Mrr-like effectors (annotated as SMEK), a newly identified group of HNH effectors, and DUF4365 effectors. Both SIR2 and TIR domains are NADases that are also found in other defence systems ^21-23^. Association of SIR2 and TIR effectors with different clades of APAZ proteins suggests their possible exchange between short pAgo systems. Direct fusions of the effector-APAZ with short pAgos are also found in the case of SIR2, TIR and DUF4365 effectors in clades S1A, S2A and S2B.

**Figure 1.**
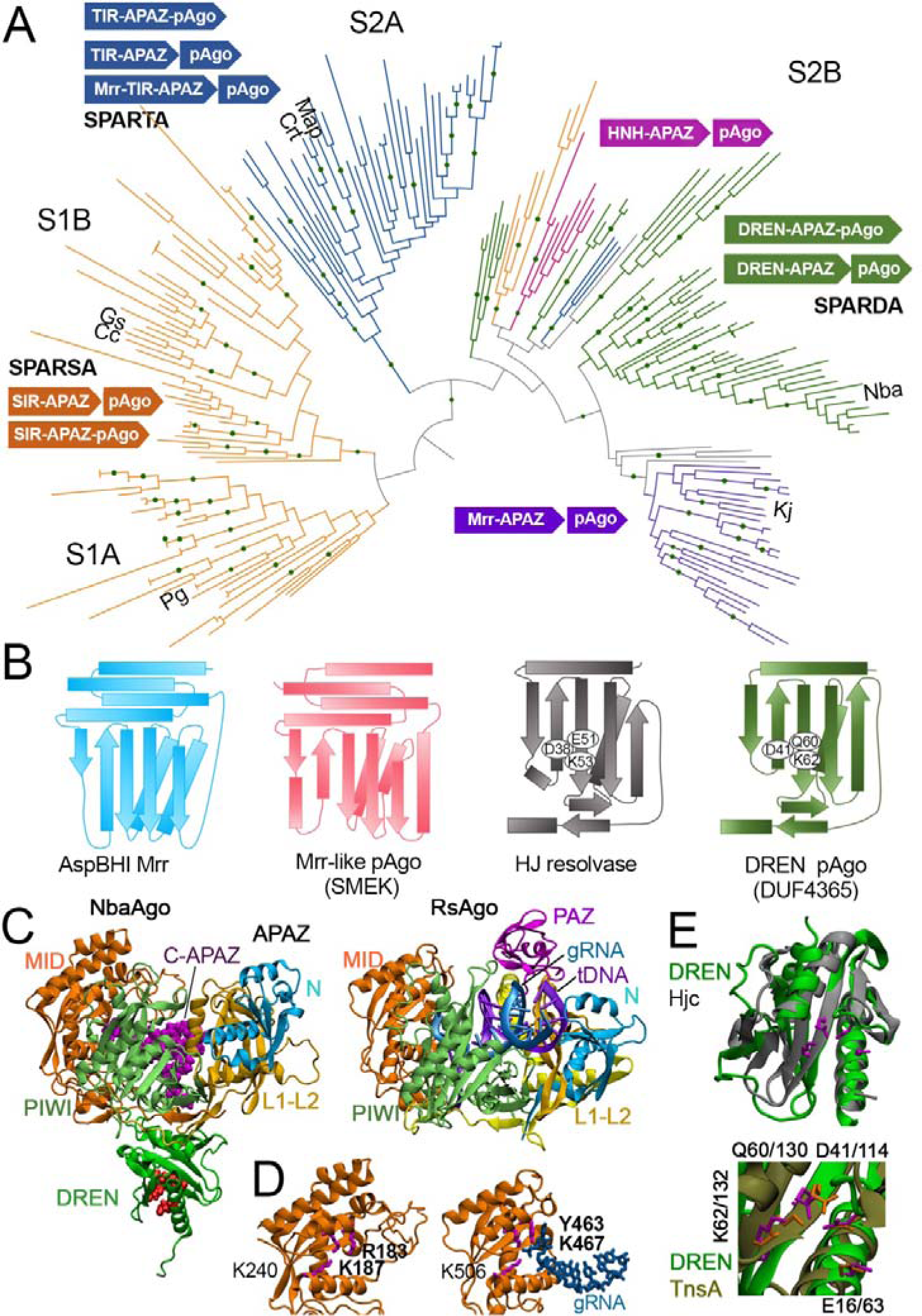
Phylogeny and structure of short pAgo systems. (A) Phylogenetic tree and operon structure of short pAgo systems. Previously studied short pAgos from the SPARSA and SPARTA systems are indicated ^18,20^. The SPARDA systems containing the DREN (DUF4365) effector are shown in green. The dots represent bootstrap values from 85% to 100%. (B) Topology of Mrr-like (SMEK) and DREN (DUF4365) domains in short pAgo systems in comparison with their closest nuclease homologues (AspBHI restriction endonuclease and Hjc resolvase). Conserved active site residues are indicated. (C) The structure of NbaSPARDA predicted using AlphaFold2 (*left*; the C-terminus of APAZ is magenta, the catalytic residues of DREN are red) in comparison with RsAgo containing bound guide RNA and target DNA (*right*; PDB: 6D8P)^29^. pAgo domains are color-coded. (D) The guide-binding pocket in the MID domain of NbaAgo (left) and RsAgo (right). Residues involved in the guide 5’-end interactions are shown in violet (the RK and YK motifs are bold). (E) Predicted structure of the DREN nuclease in comparison with the Hjc resolvase (PDB: 1GEF)^24^ and the TnsA transposase (PDB: 1T0F). The catalytic site residues are shown as stick models.

DUF4365 is the most abundant effector in the clade S2B, first found in short pAgo systems^5^. While no close homologues with known functions could be found for this domain by direct sequence search, structural modeling using the AlphaFold algorithm revealed that it belongs to the PD-(D/E)XK superfamily of metal-dependent nucleases. Mrr and HNH domains are also putative nucleases, and a short pAgo associated with an Mrr-like effector was shown to have nuclease activity *in vitro*, but was not studied in detail ^19^. In comparison with Mrr nucleases (also members of the PD-(D/E)XK superfamily), DUF4365 has a different topology, similar to Holliday junction resolving enzymes Hjc and Hje from archaea ^24,25^ (Fig. 1B). This prompted us to investigate the activities of representative DUF4365-encoding systems.

We chose short pAgo co-encoded with DUF4365-APAZ from *N. baekryungensis* (NbaAgo), an orange-colored alphaproteobacterium isolated from sea waters, for further analysis. We designated this system SPARDA, for <u>s</u>hort <u>p</u>rokaryotic <u>Ar</u>gonaute and <u>D</u>Nase/RNase-<u>A</u>PAZ. Structural modeling suggested that the two proteins form a bilobal heterodimeric complex, in which the APAZ domain is analogous to the N and L1 and L2 domains of long pAgos (Fig. 1C). The PAZ domain is missing from the structure, similarly to other short pAgo systems. The PIWI domain of NbaAgo retains a conserved RNaseH fold but lacks the catalytic tetrad residues (Extended data Fig. 1A). The MID domain forms a binding pocket for the 5’-end of a guide oligonucleotide, with an RK motif previously identified in short pAgos ^5^ (Fig. 1D, Extended data Fig. 1B). The C-terminus of DUF4365-APAZ is predicted to bind within the nucleic acid binding cleft formed by NbaAgo and APAZ, close to the guide-binding pocket of MID (Fig. 1C). The nuclease domain is connected to APAZ through a flexible linker and is modeled outside of the nucleic acid binding cleft (Fig. 1C). Its predicted structure superimposes well onto the known structure of the Hjc resolvase, while the active site and the key catalytic residues coincide with the TnsA transposase of the Tn7 family (Fig. 1E, Extended data Fig. 1C), suggesting that it is an active nuclease.

We expressed both proteins of the NbaSPARDA system in *E. coli* and purified them by metal-affinity (using a His6-tag in NbaAgo), heparin and ion-exchange chromatography. DUF4365-APAZ co-eluted with NbaAgo through all purification steps indicating that the two proteins form a stable complex, in agreement with structural predictions (Extended data Fig. 2A,B).

**Figure 2.**
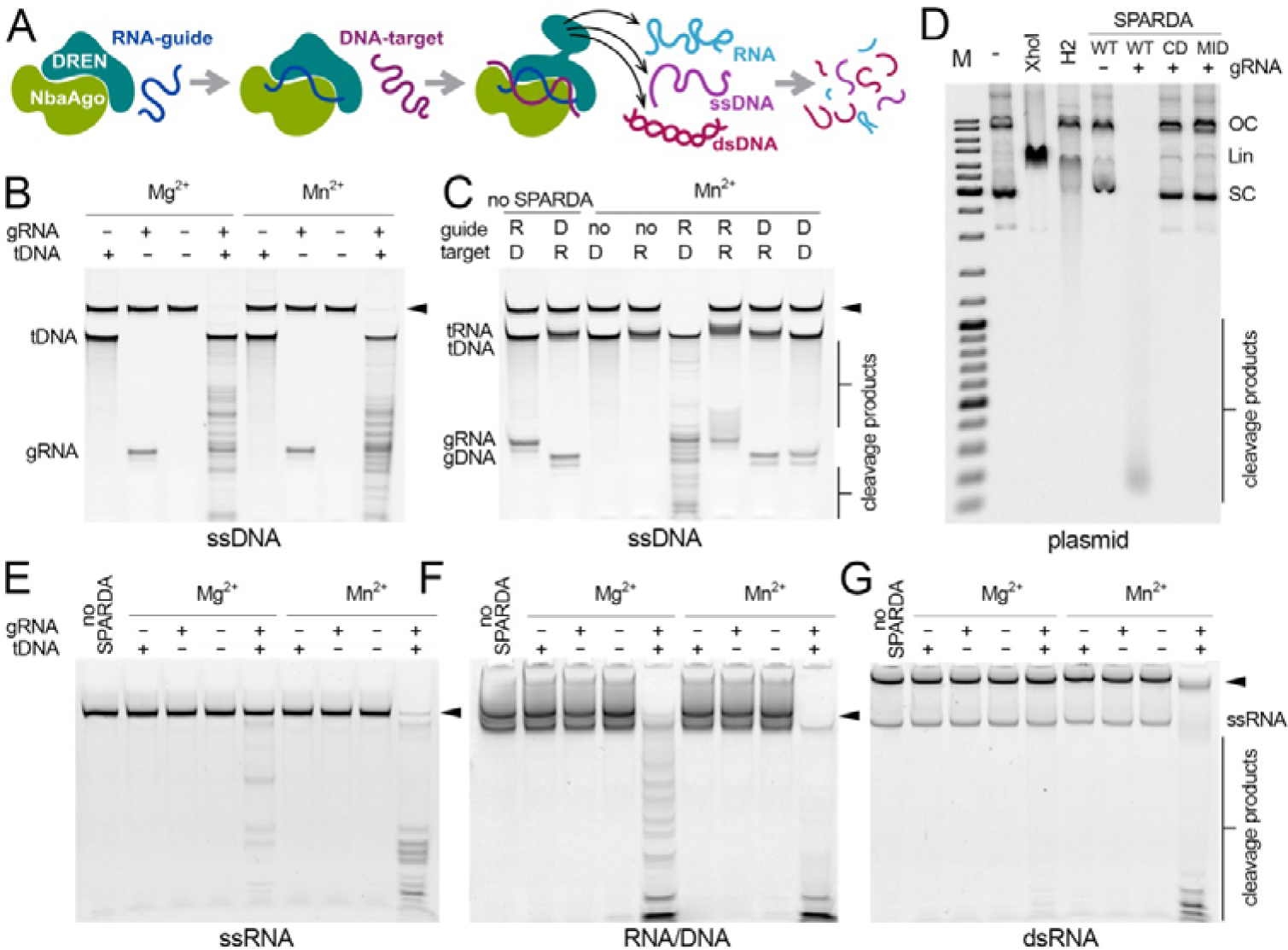
Collateral nuclease activity of SPARDA. (A) Scheme of the assay. (B) Collateral activity of SPARDA with a ssDNA substrate. (C) Analysis of the specificity of SPARDA for RNA and DNA guides and targets with a ssDNA substrate. (D) Cleavage of plasmid DNA (pJET) by SPARDA (WT, wild-type; CD, catalytically dead with substitutions in the active site of DREN; MID, substitutions in the MID domain) in the presence of gRNA and tDNA. Linear and relaxed plasmids (lanes 3 and 4) were obtained by treatment with XhoI and RNase H2. SC, supercoiled; Lin, linear; OC, open circle plasmid forms; M, length marker. (E-G) Cleavage of collateral ssRNA, RNA/DNA duplex and dsRNA, using 5’-FAM-labeled RNA, visualized by fluorescence scanning (see Fig. S5C-E for SYBR Gold staining of the same gels). Positions of collateral substrates are indicated with arrowheads; guides, targets and cleavage products are indicated. See Table S1 for oligonucleotide sequences.

### DNA targeting by SPARDA activates its effector nuclease

Since the nuclease domain in SPARDA is located outside of the guide-target binding cleft, we proposed that it may target collateral nucleic acid substrates. Previously studied short pAgos were shown to use RNA guides and preferentially recognize single-stranded DNA targets ^18,20,21^. Therefore, to analyze the potential nuclease activity of SPARDA, we incubated the purified complex with small guide RNA (gRNA), complementary target DNA (tDNA) and a collateral ssDNA substrate, bearing no complementarity to gRNA or tDNA (Fig. 2A). No cleavage of substrate DNA was observed with either gRNA or tDNA alone (Fig. 2B). In contrast, substrate DNA was fully degraded when gRNA and tDNA were present together (Fig. 2B, lanes 4 and 8). No cleavage was observed with other combinations of guide and target oligonucleotides (Fig. 2C), demonstrating that SPARDA is activated only after RNA-guided DNA target recognition. Collateral DNA cleavage was observed even with very short DNA targets (20 nt), corresponding to the length of gRNA (Extended data Fig. 3A,B). Double-stranded target DNA could not efficiently induce collateral DNA cleavage (Extended data Fig. 3C), suggesting that SPARDA cannot unwind dsDNA targets, similarly to other pAgo proteins ^15,26,27^.

**Figure 3.**
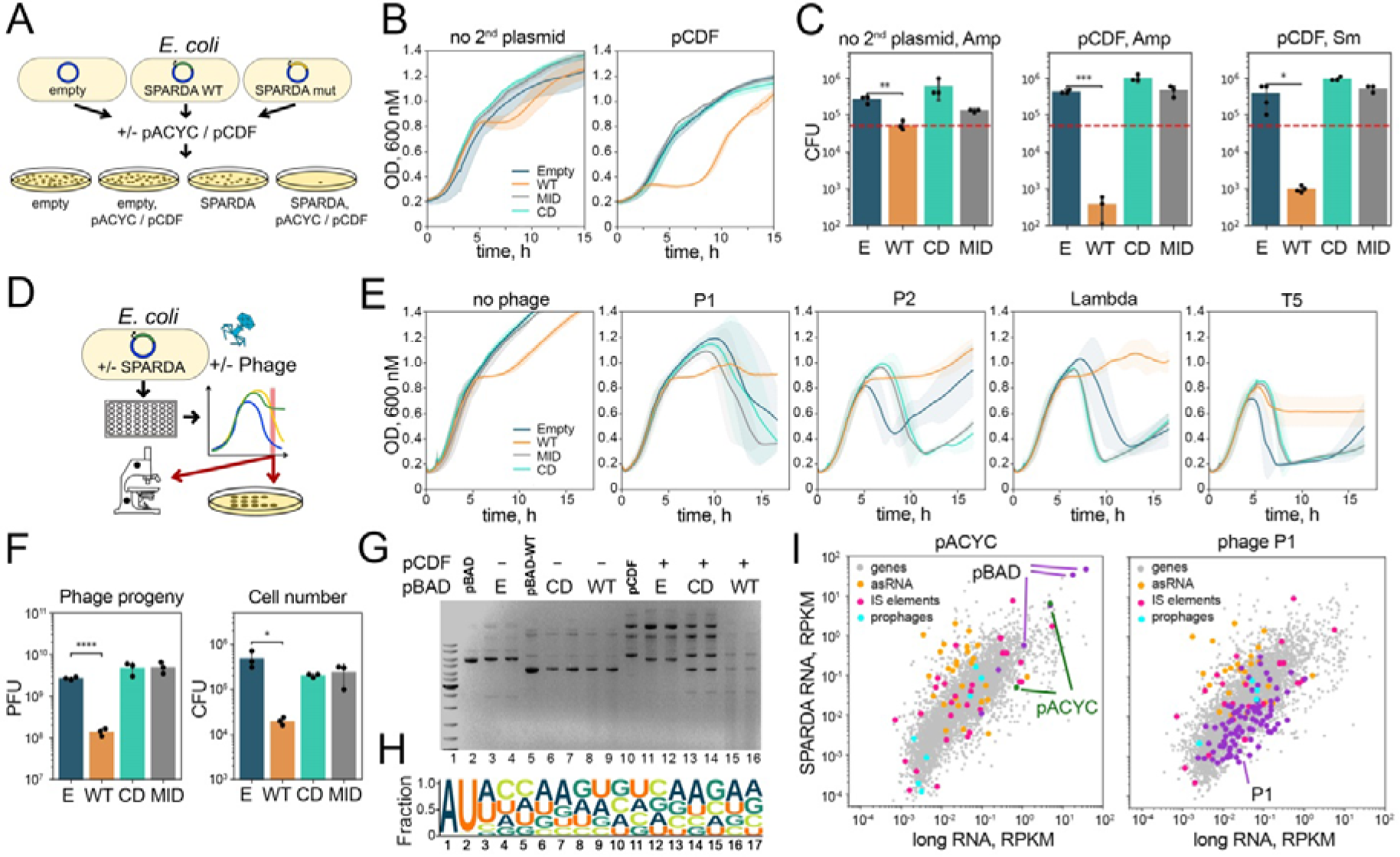
SPARDA is active against plasmids and phages. (A) Scheme of the plasmid interference assay. (B) Growth of *E. coli* strains expressing wild-type (WT) or mutant SPARDA (MID, substitutions in the MID domain; CD, catalytically dead), or containing a control empty plasmid (‘E’), in the absence and in the presence of the interfering plasmid pCDF. Means and standard deviations from 6 replicate experiments. (С) Numbers of viable cells expressing SPARDA (WT or its MID and CD mutants) or containing empty pBAD (‘E’), depending on the presence of the pCDF plasmid, at 2.5 h after SPARDA induction. Means and standard deviations from 3 or 4 replicate experiments. Statistically significant changes in CFU numbers are indicated (*p<0.05; **p<0.01, *** p<0.001). (D) Scheme of the phage assays. (E) Growth and collapse of *E. coli* cultures (MG1655) infected with phages P1, P2, T5 and λ (MOI 3.6×10^-4^ for T5 and P1, 3.6×10^-3^ for λ, 3.6×10^-6^ for P2), expressing WT or mutant SPARDA. Means and standard deviations from 5 replicate experiments. (F) Phage titers (PFU) and viable cell counts (CFU) in P1-infected cultures of *E. coli*, depending on the expression of WT SPARDA and its mutants (in BL21, analyzed at 9 hours of infection with MOI 3.6×10^-4^; Extended data Fig. 8). Means and standard deviations from 3 replicate experiments. Statistically significant changes in PFU and CFU numbers are indicated (*p<0.05; *** p<0.001). (G) Analysis of plasmid DNA integrity. Plasmid DNA was purified from *E. coli* strains containing pBAD (empty ‘E’, or encoding SPARDA, WT or CD, lanes 3-4 and 6-9), or pBAD with pCDF (lanes 11-16), at 3.5 h of growth after induction of SPARDA. For each condition, the results of two replicate experiments are shown. Lanes 2, 5 and 10 contain control samples of intact pBAD, pBAD-WT SPARDA and pCDF. (H) Nucleotide bias of gRNAs associated with SPARDA *in vivo* (from the first replicate experiment with the pACYC plasmid). (I) Correlation between the abundance of gRNAs and the RNA transcriptome of *E. coli* strains expressing SPARDA from pBAD and containing pACYC (*left*, average from three independent replicates) or infected with phage P1 (*right*, average from two independent replicates). Plasmid genes are shown in blue and violet (pBAD and pACYC); phage genes are shown in magenta. Regulatory antisense RNAs, prophage genes, and IS elements are shown in orange, pink and turquoise, respectively. The amounts of RNAs are shown in RPKM (reads per kilobase per million aligned reads in each library; for each gene, only reads corresponding to the sense DNA strand were counted).

SPARDA was active with Mg^2+^ and Mn^2+^ ions (1-10 mM) but not with Ca^2+^, Zn^2+^ or Co^2+^ or without divalent cations (Fig. 2B and Extended data Fig. 3D,E). With Mg^2+^, the highest activity was observed at low ionic strength; with Mn^2+^, DNA cleavage was activated at increased ionic strength (Extended data Fig. 3D). With both Mg^2+^ and Mn^2+^, the cleavage was efficient at pH 7.5 and 8.5, but was inhibited at pH 6.5 (Extended data Fig. 3F). With both cations, SPARDA was highly active between 25°C and 40 °C, and was inactivated above 45 °C (Extended data Fig. 3G). Therefore, SPARDA can efficiently process collateral substrates in the physiological range of conditions.

Previously studied pAgos and eAgos have distinct requirements for guide nucleic acids, including preferences for the 5’-nucleotide and certain guide length^2^. Equally high activity of SPARDA was observed with 16-24 nt gRNAs (Extended data Fig. 4A,B), indicating that varying gRNAs can be successfully accommodated within the complex. SPARDA was active with both 5’-P and 5’-OH gRNAs (Extended data Fig. 4C) but was strictly specific for the 5’-adenine in gRNA (Extended data Fig. 4D), likely because it was bound in the MID pocket, similarly to long pAgos ^28-32^. Recent analysis indicated that SPARTA and SPARSA complexes might have preferences for the second guide nucleotide *in vivo* ^18,20^. Indeed, SPARDA was active with gRNAs containing second U or C but not G or A (Extended data Fig.4E), indicating that the second guide nucleotide is also recognized specifically (Extended data Fig. 4F).

**Figure 4.**
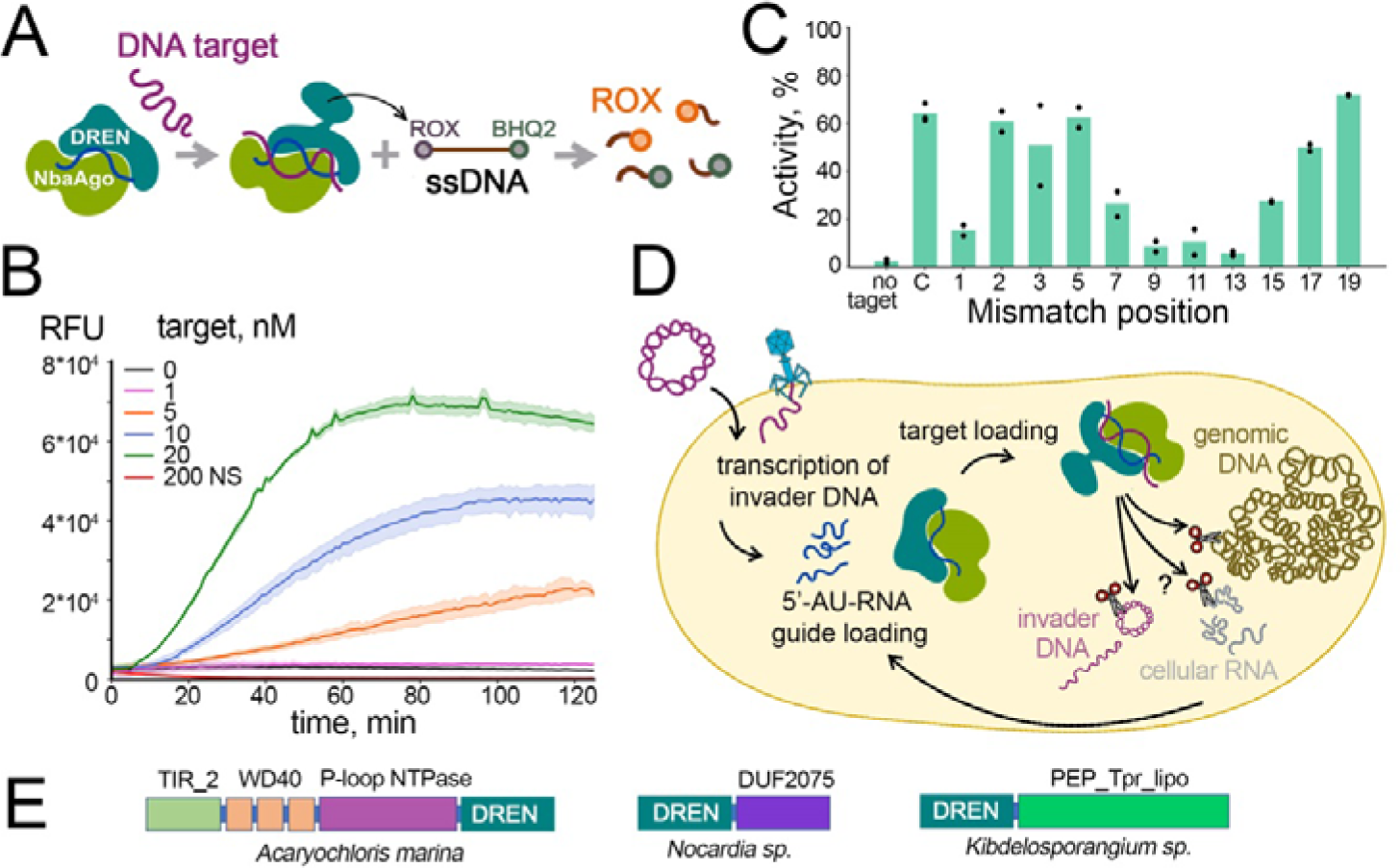
Repurposing of SPARDA for DNA detection and the proposed mechanism of SPARDA. (A) Scheme of the fluorescent beacon assay. (B) Fluorescence readout with a ssDNA beacon depending on the concentration of single-stranded tDNA (from 1 to 20 nM) or in the presence of 200 nM nonspecific DNA (NS). Means and standard deviations from three independent experiments. (C) Effects of single-nucleotide mismatches at indicated positions of tDNA on activation of SPARDA, detected by cleavage of the beacon DNA. Means from two or three independent experiments. (D) The proposed model of DNA targeting, nuclease activation and cell protection by SPARDA. (E) Examples of DREN fusions with other domains potentially serving defensive or adaptive functions (from left to right, WP_012162657.1, WP_197288423.1, WP_198151253.1). The TIR domains are found in SPARTA and Thoeris defence systems ^18,22^; the P-loop-NTPase domain together with WD40 domains can participate in signal transduction; DUF2057 is a predicted Schlafen DNA/RNA helicase with a suggested role in phage attenuation and abortive infection ^54^; PEP_Tpr_lipo is likely involved in protein export/sorting and bacterial adaptation ^55^.

Since various PD-(D/E)XK nucleases can cleave a wide spectrum of DNA and RNA substrates^33,34^, we tested whether SPARDA may degrade other collateral substrates. Wild-type SPARDA efficiently degraded circular double-stranded plasmid DNA in the presence of gRNA and tDNA, but not in control reactions without gRNA (Fig. 2D, Extended data Fig. 5A). Linear plasmid and its fragments were also completely digested within minutes (Extended data Fig. 5B). SPARDA could also process single-stranded RNA and double-stranded RNA-DNA duplex, in the presence of either Mg^2+^ or Mn^2+^ (Fig. 2E,F, Extended data Fig. 5C,D). Double-stranded RNA was cleaved incompletely, indicating that secondary structures formed in RNA transcripts may protect them from cleavage (Fig. 2G, Extended data Fig. 5E). Control reactions performed in the absence of collateral substrates demonstrated that a fraction of gRNA and tDNA was also cleaved by SPARDA, likely corresponding to unbound guides and targets (Extended data Fig. 5F).

Mutation of the RK motif (R183A/K187A/K240A) in the guide-binding pocket in the MID domain inhibited the nuclease activity of SPARDA, tested on dsDNA or ssDNA substrates (Fig. 2D, Extended data Fig. 2D), confirming that it depends on guide-directed target recognition by NbaAgo. The nuclease activity of SPARDA was fully inhibited by mutations in the predicted active site of DUF4365 (triple substitution D41A/Q60A/K62A; Fig. 2D, Extended data Fig. 2C), confirming that it is a novel type of indiscriminate nuclease, which we dubbed <u>D</u>NA/<u>R</u>NA <u>e</u>ffector <u>n</u>uclease (DREN).

### SPARDA is a heterodimer of NbaAgo and DREN-APAZ

Previously studied SPARTA systems were shown to form tetramers of pAgo-TIR-APAZ heterodimers after target DNA recognition ^18^. We analyzed the oligomeric state of the free SPARDA complex in the absence of guides and targets by size exclusion chromatography coupled with multi-angle light scattering (SEC-MALS). This analysis indicated that SPARDA is a stable heterodimeric complex with an absolute molecular weight of 104.8 kDa, very close to the predicted value of 106 kDa (54.4 kDa NbaAgo and 51.5 kDa DREN-APAZ) (Extended data Fig. 6A,B). We further analyzed possible changes in the conformation/oligomerization of SPARDA upon guide and target binding by size exclusion chromatography. Incubation of guide-free SPARDA with gRNA or gRNA-tDNA duplex slightly increased its apparent molecular weight and hydrodynamic radius (Extended data Fig. 6C), likely as a result of nucleic acid binding and conformational rearrangement of SPARDA. An additional peak likely corresponding to dimers of the heterodimeric SPARDA complex was observed in the presence of gRNA alone or gRNA and tDNA (Extended data Fig. 6C,D). Electrophoretic analysis of nucleic acids bound to SPARDA in each fraction indicated that this peak primarily contained gRNA, while both gRNA and tDNA were mainly located in the peak of the heterodimeric SPARDA complex (Extended data Fig. 6E). Therefore, SPARDA does not undergo an oligomeric state transition and hence is activated differently upon target recognition.

### SPARDA protects host cells from invaders

Previously studied CRISPR-Cas systems with collateral nuclease activity (Type III, Type V and Type VI) were proposed to prevent propagation of phages and plasmids by inducing cell death or dormancy ^35-38^. To elucidate whether SPARDA can protect the *E. coli* host from invaders, we first measured survival of cells co-transformed with a SPARDA-encoding plasmid (or a control empty plasmid, pBAD) and additional ‘interfering’ plasmids from different groups of incompatibility (Fig. 3A). The expression of SPARDA delayed cell growth (Fig. 3B) and dramatically decreased the number of viable cells (CFU, colony forming units) in the case of plasmids pACYC and pCDF (Fig. 3C, Extended data Fig. 7A,B,C). Smaller but still significant effects were also observed for other tested plasmids (Extended data Fig. 7D). Some inhibition of cell growth by SPARDA was also visible in the absence of the second plasmid (Fig. 3B,C). No decrease in cell viability was observed in the case of mutations in the guide-binding pocket in the MID domain of NbaAgo and in the active site of DREN (Fig. 3B,C). Comparable effects were observed when the cells were grown on selective antibiotics corresponding to either the interfering plasmid (Cm^R^ for pACYC or Sm^R^ for pCDF) or the SPARDA plasmid itself (Amp^R^) (Fig. 3, Extended data Fig. 7A,B,C). This suggested that SPARDA likely acts by inducing cell death or dormancy rather than by specifically targeting the interfering plasmid.

We next examined whether SPARDA can protect bacterial population from phages, by analyzing growth and collapse of *E. coli* cultures containing or lacking SPARDA during infection with phages P1, P2, T2, T4, T5, T6, T7, Qβ, N15, 121Q and λ (Fig. 3D). In the absence of phages, expression of SPARDA delayed cell growth. In contrast, it prevented the lysis of cell culture (detected as a rapid drop in cell turbidity) during infection with phages P1, P2, T5 and λ (Fig. 3E), but not other phages (Extended data Fig. 8A). Mutations in the MID domain of NbaAgo and in the active site of DREN abolished this effect for all phages (Fig. 3E).

Expression of SPARDA significantly reduced phage titers at the peak of infection, just before culture collapse was observed in the absence of SPARDA (∼20-fold decrease in the plaque forming units, PFU, for phage P1) (Fig. 3F, Extended data Fig. 8B). However, the expression of SPARDA did not increase but rather decreased the number of viable cells (CFU) in the same cultures (Fig. 3F), despite their turbidity was higher than that of the control culture. This indicated that SPARDA likely prevents phage replication and cell lysis but may also induce cell death during infection.

We directly visualized dead and live cells at the same stage of infection with propidium iodide (PI, red, penetrates cells with disrupted membranes) and SYTO 9 (green, stains only live cells in the presence of PI). A considerable number of dead (red) cells was observed among intact (green) cells in bacterial cultures expressing SPARDA in the absence of phages (Extended data Fig. 9A,B). During phage infection, wild-type SPARDA significantly increased the fraction of red cells with disrupted membranes. No dead cells were observed in the case of mutations in the MID domain of NbaAgo and in the active site of DREN, or in the absence of SPARDA (Extended data Fig. 9A,B). These results confirmed that SPARDA induces cell death during phage infection and may also be spontaneously activated in the absence of phages.

To test whether SPARDA can damage DNA *in vivo*, we purified plasmid DNA and total cellular DNA from *E. coli* cultures expressing SPARDA. Plasmid DNA was severely degraded in cells containing WT SPARDA, but not its CD mutant. At early stages of cell growth (3.5 h), this effect was observed only in the presence of the interfering plasmid (pCDF, Fig. 3G), consistent with the stronger growth phenotype of this strain (Fig. 3B). At later stages (6 h), plasmid DNA was degraded both in the absence and in the presence of the second plasmid (pCDF or pACYC; Extended data Fig. 9C). We also observed degradation of genomic DNA in cell cultures expressing SPARDA and infected with phage P1 (at 7 hr but not at 4 hr of infection; Extended data Fig. 9E). At the later stage, genomic DNA was also degraded in the absence of phages indicating that activation of SPARDA can occur spontaneously under our experimental conditions. No changes in DNA integrity were observed in cell cultures without expression of SPARDA or with mutant variants of SPARDA, CD or MID (Extended data Fig. 9E). We did not observe significant degradation of rRNA in any tested conditions, in the presence of either interefering plasmids or phage P1 (Extended data Fig. 9D,F). This indicates that activation of SPARDA causes direct damage to cellular DNA.

### SPARDA binds gRNAs produced from highly-transcribed host and invader DNA

To explore the specificity of nucleic acid targeting by NbaSPARDA *in vivo*, we expressed it in *E. coli* and purified bound nucleic acids. Wild-type SPARDA was associated with 17-22 nt gRNAs (Extended data Fig. 10A,B). The MID mutant of SPARDA did not contain any bound gRNAs, consistent with impaired gRNA binding by this mutant. In contrast, the CD mutant was loaded with gRNAs, confirming that its inactivity was not due to its inability to bind gRNA (Extended data Fig. 10B).

To determine the source of SPARDA-bound gRNAs, we sequenced gRNAs isolated from *E. coli* strains expressing SPARDA from the pBAD plasmid and containing an additional interfering plasmid (pACYC) or infected with phage P1. All gRNAs contained a 5’-AU motif, corresponding to the *in vitro* guide specificity, but did not have other nucleotide preferences (Fig. 3H). In parallel, we performed total RNA sequencing from the same strains and the same conditions. The amounts of gRNAs in general correlated with the transcript abundance (Spearman correlation coefficients of 0.85 and 0.84 for libraries obtained with pACYC and phage P1, respectively) (Fig. 3I, Extended data Fig. 11A). Long transcripts and short gRNAs from both plasmids were highly overrepresented relative to chromosomal genes, likely due to high transcription levels and high copy numbers of plasmid DNA. In contrast, phage transcripts and gRNAs were underrepresented, possibly as a result of depletion of phage-infected cells from the population (Fig. 3I, Extended data Fig. 11A).

The average ratios of gRNAs to long RNAs were comparable for chromosomal, plasmid and phage genes (and were even lower for pACYC and phage P1, Extended data Table 1), indicating no preferences of SPARDA for plasmid- or phage-derived gRNAs. To quantify the enrichment of gRNAs for certain transcripts, the amounts of gRNAs were normalized by the amounts of long RNAs for each gene or noncoding transcript. Several regions of preferential biogenesis of gRNAs were detected in plasmids and in the phage genome (Extended data Fig. 12 and 13). In the chromosome, gRNAs were enriched for noncoding antisense transcripts, but not for multicopy IS elements and prophages (Fig. 3I, Extended data Fig. 14). This indicated that NbaSPARDA proportionally binds gRNAs from highly transcribed genes and may have additional preferences for noncoding transcripts.

### SPARDA allows highly sensitive DNA detection

The activation of SPARDA by specific DNA targets prompted us to repurpose it for DNA detection. We incubated NbaSPARDA with small gRNA and complementary single-stranded tDNA and used beacon ssDNA substrates, containing a fluorophore and a quencher at its ends, for readout (Fig. 4A). We observed a robust increase in the fluorescence signal over time, corresponding to the measured kinetics of dsDNA cleavage by SPARDA. Titration experiments showed that the detection limit of tDNA was within nanomolar concentrations (Fig. 4B). We further explored the specificity of target detection by NbaSPARDA by introducing mismatches at different tDNA positions. Substitution of the first target nucleotide significantly decreased SPARDA activity (Fig. 4C), possibly indicating that this nucleotide in tDNA is recognized specifically (because it is not base-paired with gRNA). Substitutions in the seed region (positions 2, 3, 5, 7) and in the 3’-supplementary region (positions 15, 17, 19) of gRNA had only moderate or no effect on the SPARDA activity. In contrast, substitutions in the central part of gRNA (positions 9, 11, 13) strongly impaired activation of SPARDA, indicating that these positions are critical for specific target recognition (Fig. 4C). Therefore, SPARDA can detect closely related targets, similarly to PIWI-clade Ago proteins in eukaryotes^39^, but can also sense single-nucleotide mismatches at specific positions of tDNA.

## Discussion

Our data show that SPARDA is a two-component immune system consisting of a short pAgo protein and a previously unrecognized effector nuclease. Unlike long pAgo nucleases that directly destroy their targets ^10-13,15^, RNA-guided DNA targeting by SPARDA unleashes the nuclease activity of the DREN domain, leading to cleavage of collateral ssDNA, dsDNA, ssRNA and DNA-RNA substrates *in vitro* and indiscriminate degradation of cellular DNA *in vivo*. SPARDA is activated by plasmids and phages and prevents growth of cell cultures by an abortive infection mechanism (Fig. 4D). The DREN domain is also found in other genetic contexts and may therefore play more general functions in cell protection and adaptation (Fig. 4E).

In contrast to CRISPR-Cas systems that use genome-encoded gRNAs, pAgos rely on guides that are produced *de novo* from both host and invader nucleic acids^11,40,41^. The exact activation pathway of SPARDA remains to be established, including studies in native host species. Initial activation of SPARDA may occur spontaneously, or may be triggered by specific transcripts of the invading elements or by genetic stress induced by invaders, resulting in increased accessibility of small RNA guides and single-stranded DNA targets. While transcriptomic analysis revealed no obvious preferences for gRNAs derived from plasmids and phages, high transcription levels and high copy numbers of plasmid and phage DNA may be sufficient to overcome the activation threshold of SPARDA in the infected cells, similarly to SPARTA systems ^18^ and Cas13 nucleases^42^. This can be followed by explosive production of RNA and DNA fragments from invader elements as well as chromosomal DNA (Fig. 4D). The conformational changes of SPARDA during activation may include displacement of the C-terminus of DREN-APAZ from the guide-target binding cleft and rotation of the DREN domain to open its active site for substrates (as modeled in Fig. 1C; in the inactive conformation, the active site of DREN is likely buried within the complex).

SPARDA extends the list of prokaryotic defence systems that trigger suicidal cell response in response to invaders ^43,44^. Various clades of short pAgos and long inactive pAgos are co-encoded with predicted nucleases, NADases, membrane-targeting domains and other effectors, which may potentially induce abortive infection ^3-5,7,17^. Two studied short pAgo systems, SPARTA and SPARSA, trigger the NADase activity of their SIR2 and TIR effectors ^18,20,21^, while a ‘pseudo-short’ SiAgo causes membrane depolarization through its partners pAga1 and pAga2 ^45^. Besides SPARDA, CBASS and type III CRISPR-Cas systems encode RE, HNH and HEPN nuclease effectors with various spectra of DNase and RNase activities essential for their antiphage function^36,37,46-49^. Type VI Cas13 nucleases protect cells through indiscriminate cleavage of RNA after recognition of target RNA^35,38^. Type V Cas12 nucleases can degrade ssDNA and ssRNA after recognition of target dsDNA or RNA ^50,51^. Recently, a type V Cas12a2 nuclease was shown to process dsDNA, ssDNA and ssRNA substrates but, unlike SPARDA, it is activated by target RNA ^52^. SPARDA is one of the simplest among these systems with a distinct range of specificity.

SPARDA and other short pAgo systems encoding various effectors may potentially provide programmable tools for biotechnology ^53^. As a proof of principle, we have adopted the NbaSPARDA for detection of ssDNA targets using a fluorescent-reporter assay, complementary to existing Cas12- or Cas13-based approaches^38,51,52^. The sensitivity of this assay may be further increased by introducing a PCR step before detection ^18,53^. Short pAgos associated with Mrr and HNH nuclease effectors (Fig. 1A) may further expand the toolbox of defence systems activated by DNA target recognition. The low background activity of SPARDA, its physiological temperature range and independence of specific motifs in target DNA potentially allows its use in numerous applications, starting from nucleic acid detection *in vitro* to programmable elimination of bacterial or eukaryotic cells for microbiome engineering and therapy.

## Methods

### Bioinformatic analysis

The APAZ-containing proteins were searched for among the collection of bacterial and archaeal nonredundant proteins downloaded from the NCBI database in January 2023, with WP_067527439.1, WP_067969880.1, WP_033317603.1, WP_166102679.1, WP_073409929.1 and WP_034978624.1 as queries, using the deltablast 2.12.0 tool (- num_iterations=5, -evalue=0.005). A total of 20,463 protein sequences were identified, with 1,730 being unique. To annotate conserved domains in each protein, the rpsblast 2.12.0 tool was employed (-db cdd_rps, e-value threshold of 0.0001). Among the identified proteins, 386 contained the PIWI domain, corresponding to direct fusions of the APAZ domain with short pAgos^5^. Additionally, other conserved domains were observed, including SIR2 (1,123 proteins), TIR_2 (369 proteins), SMEK (175 proteins), DUF4365 (315 proteins), and Mrr (20 proteins).

To obtain the RefSeq genome ID for each protein, the Entrez Efetch tool was utilized. Subsequently, a custom script was employed to find genome neighbors for the identified proteins. The genome neighbors were then annotated for conserved domains using rpsblast, and if any of the neighboring proteins contained the PIWI domain, the proteins were annotated as short pAgo partners (Argonaute proteins associated with APAZ).

For phylogenetic analysis, the APAZ-containing proteins were clustered using a 65% sequence identity threshold by the MMseq2 software^56^, resulting in 224 representative sequences. Known short pAgo APAZ partners, including WP_046756489.1, WP_033317603.1, WP_235024190.1, WP_006053074.1, WP_010942011.1, WP_109649956.1, and WP_092459739.1, were added to the collection, resulting in a total of 231 sequences. The sequences were aligned using MAFFT v7.310 with the parameters “--maxiterate 1000 --genafpair” ^57^. The resulting alignment was then trimmed using Trimal with a gap threshold of 0.5 ^58^. The phylogenetic tree was constructed using IQ-TREE 2.0.3 with the ModelFinder algorithm, which determined the best-fit model as LG+F+R7^59^. The tree was subjected to ultrafast bootstrap analysis with 1000 replicates^60^. The resulting tree was visualized using iTol^61^.

Protein structure prediction for NbaSPARDA was performed with AlphaFold2^62^ using ColabFold^63^ (notebook AlphaFold2-mmseqs2 with default parameters in complex modeling mode), and the best model was relaxed using amber and visualized with VMD.

### Cloning, expression and purification of SPARDA

The NbaAgo protein and its neighboring protein (protein ID WP_022673743.1 and WP_033317603.1, respectively) were codon optimized using the IDT Codon Optimization Tool for expression in *E. coli* and cloned into the pBAD-HisB expression vector under the control of an inducible *araBAD* promoter, with an additional RBS motif inserted in between them, resulting in the pBAD_NbaSPARDA plasmid. Alanine substitutions in the MID domain of NbaAgo (R183A/K187A/K240A) and in the active site of DREN (D41A/Q60A/K62A) were obtained by site-directed PCR mutagenesis.

Freshly transformed *E. coli* BL21(DE3) cells were grown in the LB medium supplemented with Ampicillin (Amp, 200 µg/ml) at 37°C with shaking (200 rpm) until OD_600_=0.5. The cells were subjected to cold shock at 0°C for 30 minutes. The expression of the proteins was induced by adding arabinose (Ara) to a final concentration of 0.05%. Protein expression was performed overnight at 16°C. The cells were harvested by centrifugation and stored at −80°C. For protein purification, the cells were resuspended in a buffer containing 20 mM HEPES pH 7.5, 1 M KCl, 10 mM K_2_HPO_4_, 5% glycerol, 2 mM DTT and 2 mM PMSF and lysed by sonication at 0 °C. The resulting lysate was clarified by centrifugation for 20 min at 15,000 rpm at 4°C in a Hitachi CR22N centrifuge and loaded onto a 1 mL Ni-sepharose column (Cytiva), pre-equilibrated with a buffer containing 20 mM HEPES pH 7.5, 0.5 M KCl, and 5% glycerol. The column was washed with 10 column volumes of the equilibration buffer, followed by 10 column volumes of the same buffer supplemented with 20 mM imidazole. The protein complex was eluted using the same buffer supplemented with 300 mM imidazole. EDTA solution was added to the eluate to a final concentration of 5 mM. The eluate was diluted 5 times with the column equilibration buffer supplemented with 5 mM EDTA and further diluted 10 times with a buffer containing 20 mM HEPES pH 7.5, 5% glycerol. The diluted sample was centrifuged for 15 min at 15,000 rpm at 4°C. The supernatant was collected and loaded onto a 5 mL Heparin-sepharose column (Cytiva), pre-equilibrated with a buffer containing 20 mM HEPES pH 7.5, 40 mM KCl, and 5% glycerol. The column was washed with the same buffer, and the protein complex was eluted using a KCl gradient (from 40 to 500 mM) in the same buffer. The peak fractions containing the protein complex were pooled and diluted with buffer containing 20 mM HEPES, pH 7.5, and 5% glycerol until the concentration of KCl reached 50 mM. The samples were centrifuged at the same conditions and loaded onto a 1 ml MonoQ column, pre-equilibrated with a buffer containing 20 mM HEPES pH 7.5, 35 mM KCl, and 5% glycerol. The column was washed with 5 column volumes of the same buffer, and the protein was eluted using a KCl gradient (from 35 to 500 mM). The peak fractions containing the protein complex were pooled, analyzed by SDS-PAGE, concentrated using an Amicon 50 kDa filtering device, and supplemented with glycerol to a final concentration of 50% and DTT to a final concentration of 2 mM. The purified protein samples were stored at −20°C for further use.

### Analysis of *in vitro* activities of SPARDA

The nuclease activity of SPARDA was analyzed using synthetic oligonucleotides, which served as the guide, target, and substrate (see Supplementary Table S1). The SPARDA complex (500 nM final concentration) was preincubated with guide RNA (200 nM) for 5 minutes at 30°C in a buffer containing 20 mM HEPES pH 7.5, 5 μg/ml BSA, 5% glycerol, 0.4 mM DTT, appropriate divalent cations and KCl as specified in the figures (5 mM Mg^2+^ and 5 mM KCl or 5 mM Mn^2+^ and 100 mM KCl in most reactions). Subsequently, 200 nM target DNA was added and the samples were incubated for additional 5 minutes. Collateral substrate oligonucleotides were added to the 100 nM final concentration (ssDNA, dsDNA, ssRNA, RNA/DNA hybrid or dsRNA, as indicated in the figures). Supercoiled plasmid DNA (pBAD-HisB) was obtained by the GeneJET Plasmid Miniprep Kit (Thermo Fisher Scientific) and was added to the 1-2 nM concentration. Following the addition of the substrates, either kinetic or endpoint experiments were conducted. In most experiments, the reactions were performed for 30 minutes at 30 °C, unless otherwise indicated. To reactions were stopped with a loading buffer containing 8 M UREA, 20 mM EDTA, 2^x^TBE. The reaction products were separated by 19% denaturing urea PAGE and visualized by SYBR GOLD staining. The gels were scanned using a Typhoon FLA 9500 scanner (GE Healthcare).

### Size-exclusion chromatography and SEC-MALS analysis

The absolute molecular mass of the SPARDA complex was determined for the WT and MID mutant NbaAgo forms devoid of nucleic acids (the OD_260_/OD_280_ ratio of 0.57-0.6) by size-exclusion chromatography (SEC) coupled to multi-angle light scattering (MALS). To this end, the NbaAgo samples (3 mg/ml in 33 μl) were loaded onto a Superdex 200 Increase 10/300 column (GE Healthcare, USA) operated on a ProStar335 system and connected to a tandem of Prostar 335 detector (Varian, Australia) and miniDAWN detector (Wyatt Technology, USA) at a constant flow rate of 1 ml/min. The column was pre-equilibrated and run with filtered (0.1 μm, Millipore) and degassed buffer containing 20 mM HEPES-KOH, pH 7.5, 100 mM KCl, 5% glycerol and 5 mM MgCl_2_. The data were analyzed in ASTRA 8.0 (Wyatt Technology, USA) using dn/dc = 0.185 and the protein extinction coefficient ε (0.1%) at 280 nm of 1.29 (mg/ml)^-1^cm^-1^. The monodispersity for a chosen peak was determined as a Mw/Mn ratio. The miniDAWN and UV detectors were also used to confirm the oligomeric nature of the minor, lower elution volume peak of the SPARDA complex with the FAM-labeled guide RNA, via the principle of static light scattering coherence of the oligomeric species (the scattered light from the dimer is twice as intense as the total light scattered from two independent monomers) ^64,65^.

Size-exclusion chromatography was used to verify the direct binding of NbaAgo with the synthetic gRNA and tDNA. Elution profiles were registered by absorbance at 280 and 500 nm upon loading of 33 μl samples (2-3 mg/ml load protein concentration) onto the column operated on a ProStar335 system (Varian, Australia) at a flow rate of 1 ml/min in the same buffer. The 500 nm absorbance track was optionally present to track the elution profile of the FAM-labeled guide in the samples. Contents of the fractions collected along each elution profile were assessed by protein or oligonucleotide polyacrylamide gel electrophoreses. The apparent molecular weight corresponding to the chromatographic peaks was determined via column calibration by protein standards (α-lactalbumin monomer 15 kDa, ovalbumin monomer 43 kDa, BSA monomer 66 kDa, BSA dimer 132 kDa, BSA trimer 198 kDa).

### Analysis of cell viability during plasmid transformation

*E. coli* BL21(DE3) cells were transformed with various plasmids, including the pBAD-HisB empty vector, pBAD_NbaSPARDA (WT, MID or CD mutants), as well as plasmids pACYC184, pCDF-duet, pMB9, and RSF1010. An overnight culture was prepared using two antibiotics (Amp and the second antibiotic corresponding to each of the plasmids) and glucose (Glc, 0.5%). The culture was then diluted 76 times in LB medium supplemented with Amp and 0.1% Ara. Aliquots of 200 µl were dispensed into a 48-well plate. For the culture growth curves, the plates were incubated for 20 hours at 30°C with shaking at 200 rpm in a CLARIOSTAR plate reader, with OD_600_ measurements every 10 min. For CFU measurements, the plate was incubated for 2.5 hours at 200 rpm at 30°C. Serial dilutions of each culture were then plated on a Petri dish containing appropriate antibiotics and 0.5% Glc (to suppress expression of SPARDA). The Petri dishes were incubated at 37°C overnight, and the number of colonies was counted the following day.

### Analysis of phage infection

The anti-phage activity of NbaSPARDA and its mutant variants was tested with different phages, including P1, P2, T2, T4, T5, T6, T7, Qβ, N15, 121Q and λ (from our laboratory collection and kindly provided by Artem Isaev and Konstantin Severinov). Phage lysates were prepared using the *E. coli* MG1655 strain (or *E. coli* DH5α Turbo, New England BiolabsNEB) in the case of N15 and Qβ phages). Phage titers were determined by counting plaque forming units (PFU). For analysis of NbaSPARDA, *E. coli* MG1655 Z1, *E. coli* BL21 (DE3) or *E. coli* DH5α Turbo and were transformed with pBAD plasmids encoding WT or mutant variants of SPARDA, or control empty pBAD, grown overnight in LB supplemented with Amp (100 μg/ml) and Glc (0.5%, to suppress premature expression of SPARDA), diluted two-fold with 50% glycerol, aliquoted, frozen in liquid nitrogen and stored at −80°C. For analysis of the growth curves, overnight bacterial cultures were obtained from frozen aliquots grown in LB with Amp and Glc and 5 μl were inoculated into 100 μl of fresh LB supplemented with CaCl_2_ (5 mM), MgSO_4_ (10 mM), Amp and Ara (0.05%) in 96-well plates. After infection with various phages, the plates were incubated at 200 rpm at 30°C in a CLARIOSTAR microplate reader and cell density was monitored by measuring OD_600_ every 10 min (Extended data Fig. 8A). Phages that were affected by the SPARDA expression were taken into further analysis. To confirm the anti-phage activity of SPARDA against P1, P2, T5 and λ, overnight bacterial cultures were obtained from frozen aliquoted cultures as described above, 10 μl were inoculated into 1 ml of LB supplemented with CaCl_2_ (5 mM), MgSO_4_ (10 mM), Amp and Ara in 24-well plates, the pages were added at the multiplicity of infection (MOI) of 3.63×10^-4^ for T5 and P1, 3.6×10^-3^ for λ and 3.6×10^-6^ for P2, and culture growth was monitored as described above (Fig. 3E).

To investigate the production of virus particles in cultures expressing SPARDA, overnight cultures of *E. coli* BL21(DE3) were inoculated into fresh medium in 24-well plates as described above and infected with P1 at MOI of 3.63×10^-4^ (Extended data Fig. 8B). To determine phage titers, 100 μl aliquots were taken from the cultures at 10 hours after phage inoculation and treated with 100 μl of chloroform for 15 s with vortexing. Serial dilutions were prepared in 10 mM MgSO_4_, 5 mM CaCl_2_ and aliquots were plated on LB plates covered with 0.3% top agar containing 10 mM MgSO_4_, 5 mM CaCl_2,_ 5% glycerol and MG1655 Z1 cells grown until OD_600_ of 0.2. Phage plaques were counted after overnight growth at 30°C. CFU numbers were determined as described above for plasmid transformation, except for the cells were washed with LB containing citrate (15 mM) and Glc (1%) prior to serial dilutions in LB, supplemented with Amp (100 µg/ml), Glc (1%) and citrate (5 mM).

### Live-dead staining of *E. coli*

The samples for live-dead staining were taken at 14 hours post infection in the same conditions as for growth curve experiments with phage P1 (Fig. 3E), using *E. coli* MG1655. The cells were rinsed twice with 1 mL 0.9% NaCl, resuspended in 0.9% NaCl to OD_600_∼0.8 and then stained with 1 mM SYTO 9 (excitation/emission maxima: 485/500 nm) and 6 mM propidium iodide (PI) (excitation/emission maxima: 490/635). Control samples with killed cells were prepared as described in the LIVE/DEAD BacLight Bacterial Viability kit L7012 (Molecular Probes). The samples were imaged with a ZEISS LSM 900 confocal laser scanning microscope (Carl Zeiss), using excitation wavelengths of 488 nm (SYTO 9) and 561 nm (PI). Cell images were taken with a 63x oil immersion objective. Contrast/brightness settings were adjusted individually for each channel using the ZEN Microscopy software (Carl Zeiss). For calculation of the numbers of red and green cells in samples expressing SPARDA infected and uninfected with phage P1, 368 and 245 cells from 15 fields of views from the respective samples were taken into analysis. Cells emitting both red and green colors were considered red (dead cells).

### Analysis of DNA and RNA degradation *in vivo*

Samples for analysis of plasmid DNA degradation were collected from *E. coli* BL21(DE3) cultures transformed with pBAD_NbaSPARDA and indicated plasmids (pACYC184 or pCDF) and grown in the same conditions as for analysis of growth curves (Fig. 3B) after 3.5 or 6 hours after induction of SPARDA. Plasmid DNA was purified using Zyppy™ Plasmid Miniprep Kit (Zymo Research) and analyzed by 1% (TAE 1^x^) agarose gel electrophoresis, followed by ethidium bromide staining. Total RNA was purified from the same cultures after 7 hours of growth using GeneJET RNA Purification Kit (Thermo Fisher Scientific) and analyzed in the same way. Samples for analysis genomic DNA and total RNA degradation during phage P1 infection were collected from *E. coli* MG1655 cultures grown in the same conditions as for analysis of growth curves with phage P1 (Fig. 3E) at 2.5, 4, 6 or 7 hours after induction of SPARDA, as indicated in the figures. The samples were centrifuged at 10,000 g for 3□min. The pellets were frozen in liquid nitrogen and stored −80□°C until further processing. Genomic DNA was extracted using GenElute™ Bacterial Genomic DNA Kit (Sigma-Aldrich) following manufacturer instructions. Total RNA was extracted using GeneJET RNA Purification Kit (Thermo Scientific) following manufacturer instructions. In case of DNA a total of 1 µg of each sample was combined with Gel Loading Dye, Purple(6X) (New England Biolabs) and loaded onto a 0.8% agarose 0.5xTBE gel supplemented with ethidium bromide. For total RNA visualization a total of 1□µg from each sample was combined with the same loading dye, heated to 70□°C for 3□min and immediately chilled on ice for 2□min. The denatured samples were loaded onto 1.2% agarose 1xTAE gel. The gels were run for 1 hour at 120□V and imaged.

### Isolation of total RNA and SPARDA-associated gRNAs

Overnight cultures (1.4 ml) of *E. coli* BL21(DE3) transformed with pBAD_NbaSPARDA (two replicates) or pBAD_NbaSPARDA and pACYC184 (three replicates) were inoculated into 0.7 L of the LB medium. The medium was supplemented with Amp (100 µg/ml), 10 mM MgCl_2_, 5 mM CaCl_2_, and 0.05% arabinose. The cultures without pACYC184 were infected with 280 µl of P1 phage with a titer of 10^6^ PFU/ml. The flasks were incubated at 30°C with shaking for 5 hours. Ten ml was taken from each flask, pelleted by centrifugation, washed twice with ice-cold TNE buffer, and aliquoted. The aliquots were used for total RNA purification using the JeneGet RNA purification kit (Thermo Fisher Scientific). The remaining culture was precipitated, washed once with LB and stored at −80°C. The next day, the cells were resuspended in 15 ml of ice-cold lysis buffer (HEPES, pH 7.5, 1 M NaCl, 10 mM KH_2_PO_4_, 2 mM PMSF, 5 mM β-mercaptoethanol) and lysed by sonication. The lysates was clarified by centrifugation as described above and 200 µl of TALON resin suspension (Takara Bio), equilibrated with the same buffer, was added. The samples were incubated with rotation at 4°C for 3 hours. After incubation, the resin was precipitated at 1000 rpm at 4°C in an Eppendorf centrifuge, and the supernatant was discarded. The resin was washed once with 5 ml of ice-cold buffer containing 30 mM HEPES, pH 7.5, 0.5 M KCl, 10 mM KH_2_PO_4_, 5% glycerol), once with 1 ml of the same buffer, and once with the same buffer supplemented with 5 mM imidazole. The SPARDA complex was eluted with 0.5 ml of the same buffer supplemented with 300 mM imidazole. 200 µl of the eluate were used for nucleic acid purification by phenol-chloroform extraction. 100 µl of equilibrated phenol and 100 µl of chloroform were added, the samples were intensively shaken and centrifuged. This procedure was repeated, and nucleic acids were precipitated by adding 20 µl of 3M ammonium acetate, 0.8 ml of 96% ethanol, and 1 µl of RNase-free glycogen. After incubation at −20°C overnight, nucleic acids were precipitated by centrifugation at 13,000 rpm at 4°C in an Eppendorf centrifuge for 1 hour, washed with 0.7 ml of ice-cold 80% ethanol, dried, and dissolved in mQ-grade water. Total RNA and short RNAs purified from the SPARDA complex were processed with RQ DNAse (Qiagen) following the manufacturer’s protocol and subjected to phenol-chloroform extraction, as described above.

### Small RNA and total RNA sequencing

Ribosomal RNA was depleted from total RNA preparations with the NEBNext rRNA Depletion Kit (Bacteria) (New England Biolabs). Total RNA libraries were prepared with NEBNext Ultra II Directional RNA Library Prep Kit for Illumina. Libraries of small RNAs were prepared according to the previously published splinted ligation protocol ^41^. Briefly, nucleic acids extracted from SPARDA were phosphorylated by T4 polynucleotyide kinase (NEB) according to the manufacturer’s protocol and were separated by 18% PAGE, small RNAs (14-24 nt) were eluted from a crushed piece of the gel in 0.4 M NaCl overnight at 21 °C, ethanol precipitated, dissolved in water, phosphorylated with polynucleotide kinase (New England Biolabs). Libraries were prepared with NEBNext® Small RNA Library Prep Set for Illumina® (Multiplex Compatible). All RNA libraries were sequenced using the HiSeq2500 platform (Illumina) in the rapid run mode (50-nucleotide single-end reads). The list of sequenced RNQA libraries is presented in Supplementary Table S2.

### Analysis of high-throughput sequencing data

Raw reads were quality checked with FastQC (v. 0.12.1), adaptors were trimmed and reads with the length less than 14 nt were discarded from further analysis with Trimmomatic (v. 0.39). Reads were aligned to the reference genomic and plasmid DNA (Refseq accession number NC_012971.2 for BL21(DE3), GeneBank number OP279344 for phage P1) allowing no mismatches for short reads or 1 mismatch for long reads, using Bowtie (v. 1.0.0). Nucleotide logo was calculated and visualized using logomaker package for python ^66^. For counting reads in genome features total RNA sequences were reverse complemented with seqtk program and all libraries were aligned to the bacterial chromosome, phage or plasmid DNA allowing no mismatch for short reads and one mismatch for long reads. For chromosomal mapping, the MG1655 genome was as a reference (RefSeq accession number NC_000913.3), because it contains functional annotations of various groups of genes (including noncoding and antisense regulatory RNAs). The number of reads in genomic features was calculated by htseq-count (v. 0.13.5) in strand specific and ‘union’ mode (-s yes -m union). The resulting numbers of reads were normalized by gene length and by the total numbers of mapped reads and visualized with custom Python scripts (https://github.com/prostovna/short-argonaute-guides.git).

### Fluorescence assay of SPARDA activity

For fluorescence assay, the following synthetic oligonucleotides were used: 20 nt gRNA, 20 nt tDNA, and BHQ2-DNA-32nt-ROX or DNA-32nt-ROX as substrates (see Supplementary Table S1). The SPARDA complex (2 µM) was preincubated with gRNA (700 nM) for 15 minutes at 30°C and then cooled on ice. Subsequently, tDNA (700 nM) and ssDNA substrate (700 nM) were added, the samples were aliquoted into a 384-well plate on ice, and the plate was placed in a CLARIOSTAR reader operating in enhanced dynamic range mode. Measurements were taken every 1 minute at 30°C for 2 hours.

### Statistical analysis

For statistical analysis of data of plasmid and phage experiments, the statannot Python package was used. P-values were calculated by applying t-test for independent samples. For Spearman correlation analysis, pandas package method corr was used.

## Acknowledgements

We thank Alexei Aravin for inspiration of this work and helpful discussions, Anna Olina for help in analysis of SPARDA-associated small RNAs, Mayya Petrova for plasmids, Artem Isaev and Konstantin Severinov for phages, Svetlana Pitelyak for help with phage assays and Nikita Mikhaylov for invaluable programming clues. This work was supported by the Russian Science Foundation (grant 22-14-00182 to A.Kulbachinskiy).

## Data and Code Availability

The data generated during this study are included in the published article, Extended Data and Supplementary data. The gRNA and total RNA sequencing datasets generated in this study are available from the Sequence Read Archive (SRA) database (BioProject ID PRJNA975910). The code used for data analysis is available at the GitHub repository https://github.com/prostovna/short-argonaute-guides.git. All primary data are available from the corresponding authors upon request.

## Author contributions

Conceptualization and supervision: M.P., D.E. and A.Kulbachinskiy. Phylogenetic and structural analysis: M.P. Cloning and purification of SPARDA: A.Kanevskaya and M.P. Experiments *in vitro*: A.Kanevskaya under supervision of M.P. Experiments *in vivo*: A.Kanevskaya, V.P., M.P. SEC-MALS analysis: N.N.S., K.T. and A.Kanevskaya. Preparation and sequencing of RNA libraries: A.Kanevskaya, L.Lisitskaya and M.P. Analysis of sequencing data: M.P. Writing - original draft: A.Kulbachinskiy. Writing - review and editing: A.Kulbachinskiy and M.P with contribution from all of the authors.

## Additional information

**Supplementary information.** The online version contains supplementary material.

**Correspondence and requests for materials** should be addressed to M.P. or A.Kulbachinskiy.

## Extended Data Figures

**Extended Data Figure 1.**
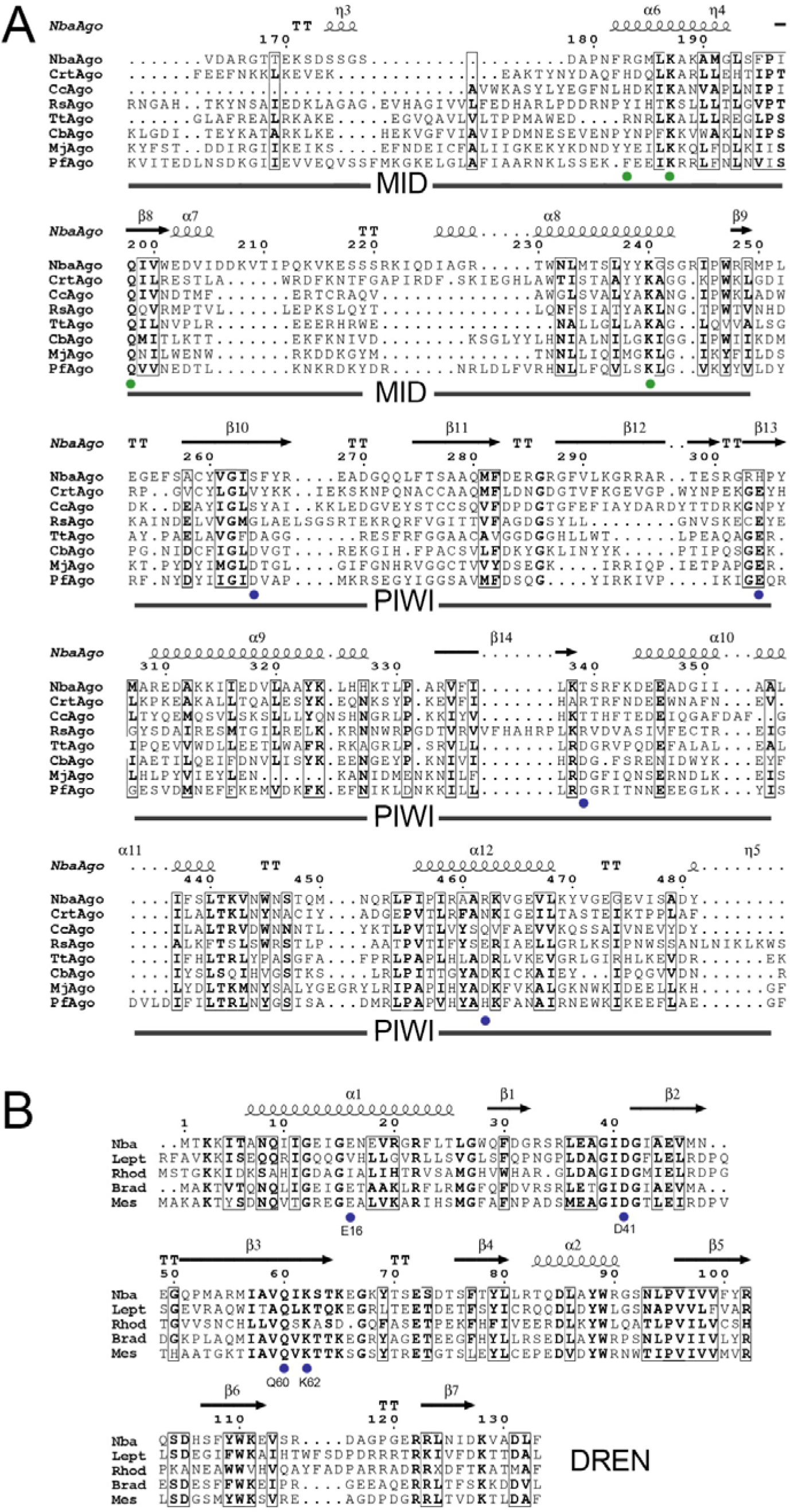
Conservation of key structural elements in SPARDA. (A) Alignment of the MID and PIWI domains in short pAgos from the SPARDA (NbaAgo), SPARTA (CrtAgo), SPARSA (CcAgo) systems and in long pAgos (RsAgo, TtAgo, CbAgo, MjAgo, PfAgo). Residues involved in interactions with the guide 5’-end in the MID pocket are shown with green dots (YKQK in RsAgo, RKQK in NbaAgo, see Fig. 1D). The active site residues in the PIWI domain in active long pAgo nucleases (TtAgo, CbAgo, MjAgo, PfAgo) are shown with blue dots. The source of pAgo proteins: NbaAgo - *Novosphingopyxis baekryungensis* (WP_022673743.1), CrtAgo – *Thermoflavifilum thermophilum* (WP_092459739.1), CcAgo - *Caballeronia cordobensis* (WP_053571899.1), RsAgo - *Rhodobacter sphaeroides* (WP_026824436.1), TtAgo - *Thermus thermophilus* (WP_011174533.1), CbAgo - *Clostridium butyricum* (WP_058142162.1), MjAgo - *Methanocaldococcus jannaschii* (WP_010870838.1), PfAgo - *Pyrococcus furiosus* (WP_011011654.1). The amino acid numbering is shown for NbaAgo. (B) Residues of the nuclease active site in DUF4365 in SPARDA systems from various species. Nba - *Novosphingopyxis baekryungensis* (WP_033317603.1), Lept - *Leptolyngbya sp*. (WP_006512593.1), Rhod - *Rhodoplanes elegans* (WP_111355734.1), Brad - *Bradyrhizobium sp*. (WP_008137934.1), Mes - *Mesorhizobium sp*. (WP_136091384.1). The predicted active site residues are indicated with blue dots. The amino acid numbering is shown for NbaDUF4365. The predicted elements of the secondary structure are indicated (α-helices, π-helices and β-strands; strict β-turns are shown as TT letters).The alignment is generated with ESPript 3.0.

**Extended Data Figure 2.**
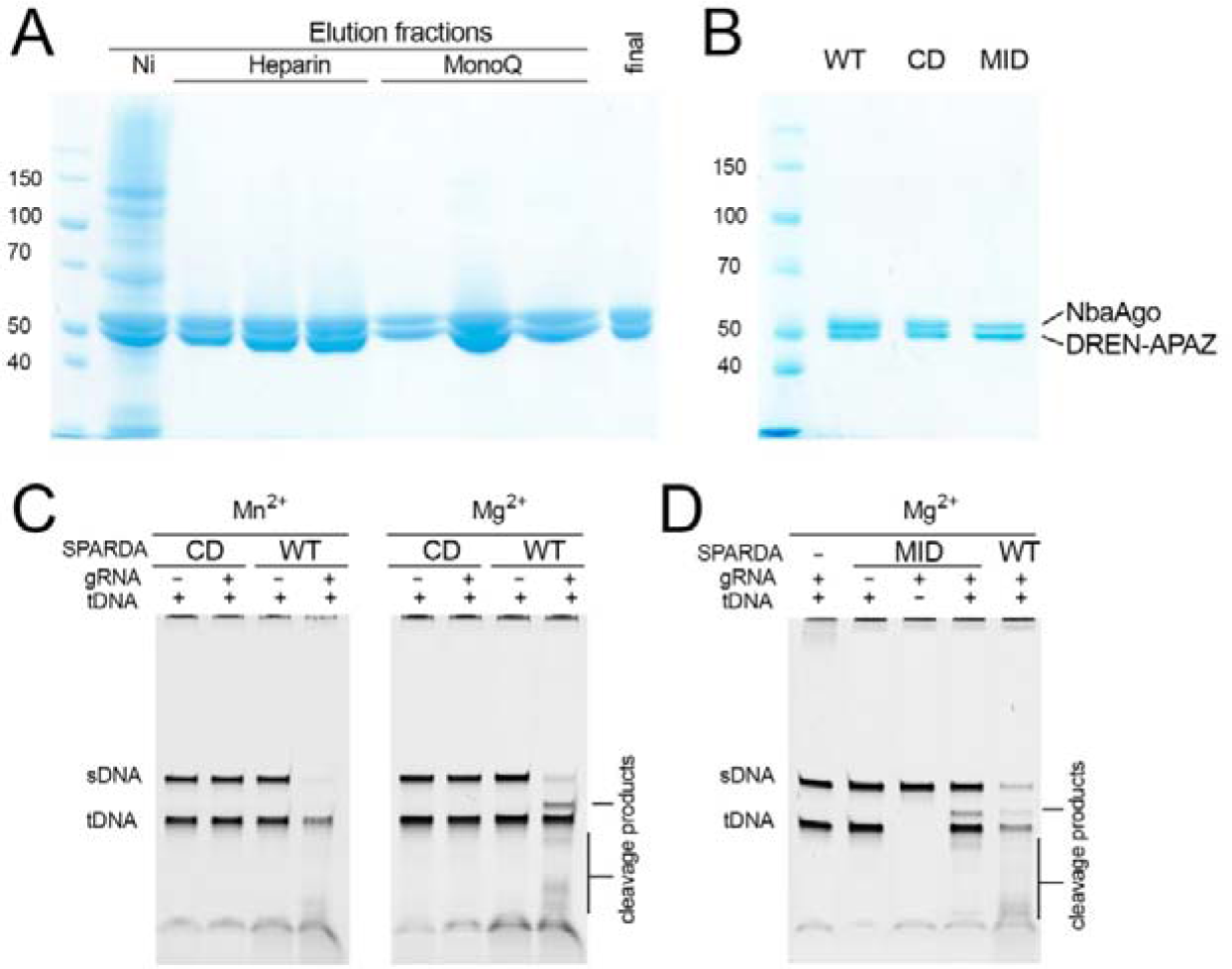
Purification of wild-type and mutant SPARDA complexes. (A) Co-elution of NbaAgo and DREN-APAZ during Ni-chelating, heparin and anion-exchange (MonoQ column) chromatography steps. Individual fractions and the final protein sample are shown for the wild-type complex. (B) Final purified samples of wild-type (WT) SPARDA and its mutants with substitutions in the active site of DREN (CD, catalytically dead) and in the MID pocket of NbaAgo (MID). (C) Comparison of the activity of WT and CD variants of SPARDA with gRNA (20 nt 5’-A-RNA), tDNA and single-stranded collateral single-stranded substrate DNA (sDNA) in the presence of Mg^2+^ and Mn^2+^. (D) Comparison of the activity of WT and MID variants of SPARDA with gRNA (20 nt 5’-A-RNA), tDNA and single-stranded sDNA in the presence of Mg^2+^. The experiments in C and D were performed with WT SPARDA samples containing admixture of endogenous gRNAs (see Extended Data Fig. 10). Positions of tDNA, sDNA and cleavage products are indicated.

**Extended Data Figure 3.**
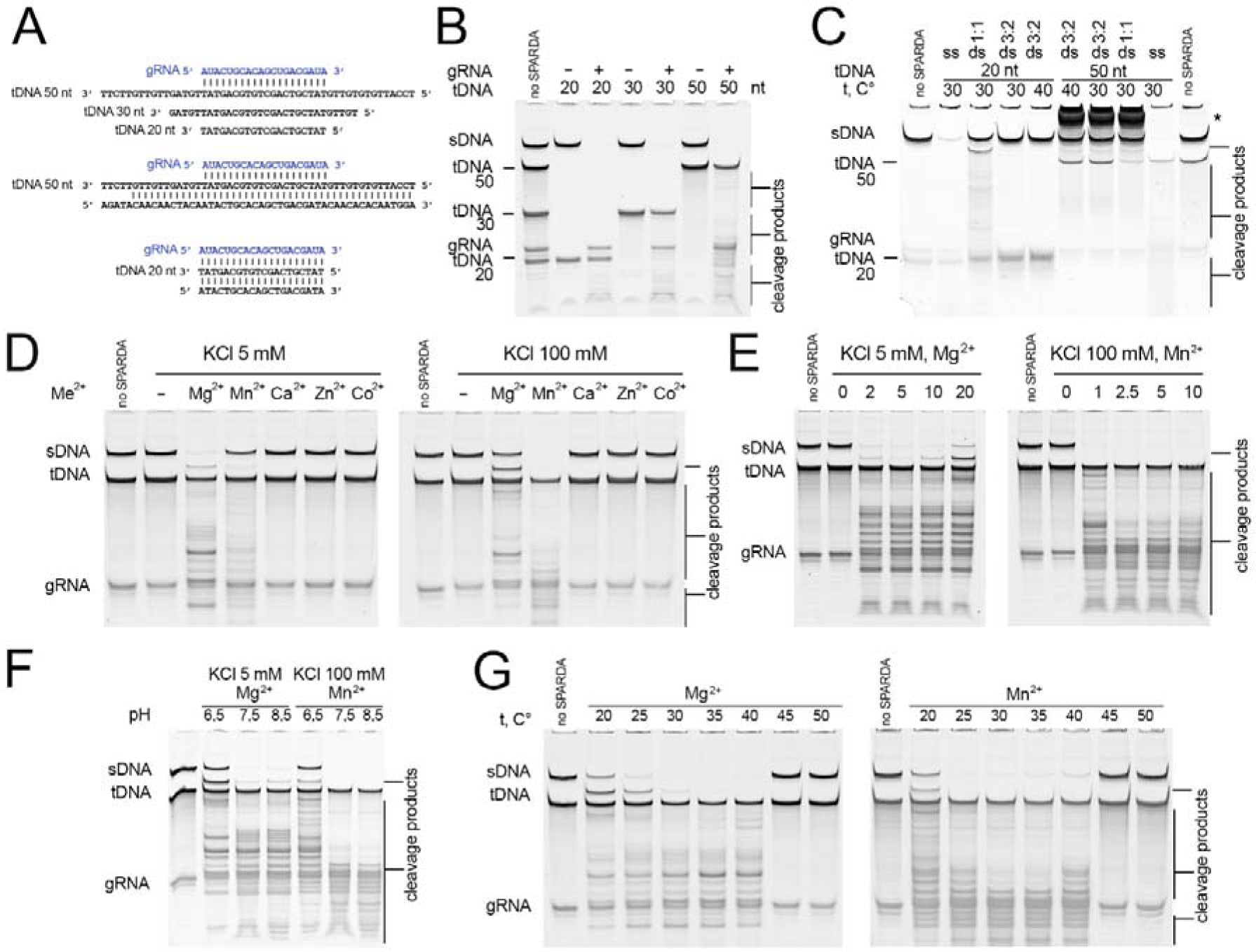
Analysis of the optimal conditions for SPARDA activity *in vitro*. (A) Structure of gRNA and tDNAs (single-stranded or double-stranded) used in the assays. (B) Activity of SPARDA with single-stranded tDNAs of various lengths (20, 30, and 50 nt), in the absence and in the presence of gRNA. (C) Activity of SPARDA with double-stranded tDNA (20 nt, lanes 3-5, or 50 nt, lanes 6-8), tested for 60 minutes at 30 °C or 40 °C. Control reactions were performed with single-stranded tDNA (lanes 2 and 9). The non-target DNA strand was annealed with the complementary target strand at the 1:1 or 3:2 ratio. While low level of SPARDA activity was observed at the 1:1 ratio (lanes 3 and 8), it was suppressed at the 3:2 ratio (lanes 4-5 and 6-7), indicating that this activation resulted from the presence of non-annealed single-stranded tDNA. The position of non-denatured 50 nt duplex DNA is indicated with an asterisk. The reactions in B and C were performed in the presence of Mn^2+^. (D) Testing of the SPARDA activity with single-stranded tDNA in buffers containing various divalent cations (5 mM each), with either 5 mM or 100 mM KCl. (E) Activity of SPARDA with different concentrations of Mg^2+^ (left, with 5 mM KCl) or Mn^2+^ (right, with 100 mM KCl). (F) Activity of SPARDA at various pH with Mg^2+^ (left, 5 mM KCl) or Mn^2+^ (right, 100 mM KCl). (G) Activity of SPARDA at various temperatures with Mg^2+^ (left, 5 mM KCl) or Mn^2+^ (right, 100 mM KCl). All reactions except for panels C and G were performed at 30 °C. Positions of gRNA, tDNA, single-stranded collateral substrate DNA (sDNA) and the cleavage products are indicated.

**Extended Data Figure 4.**
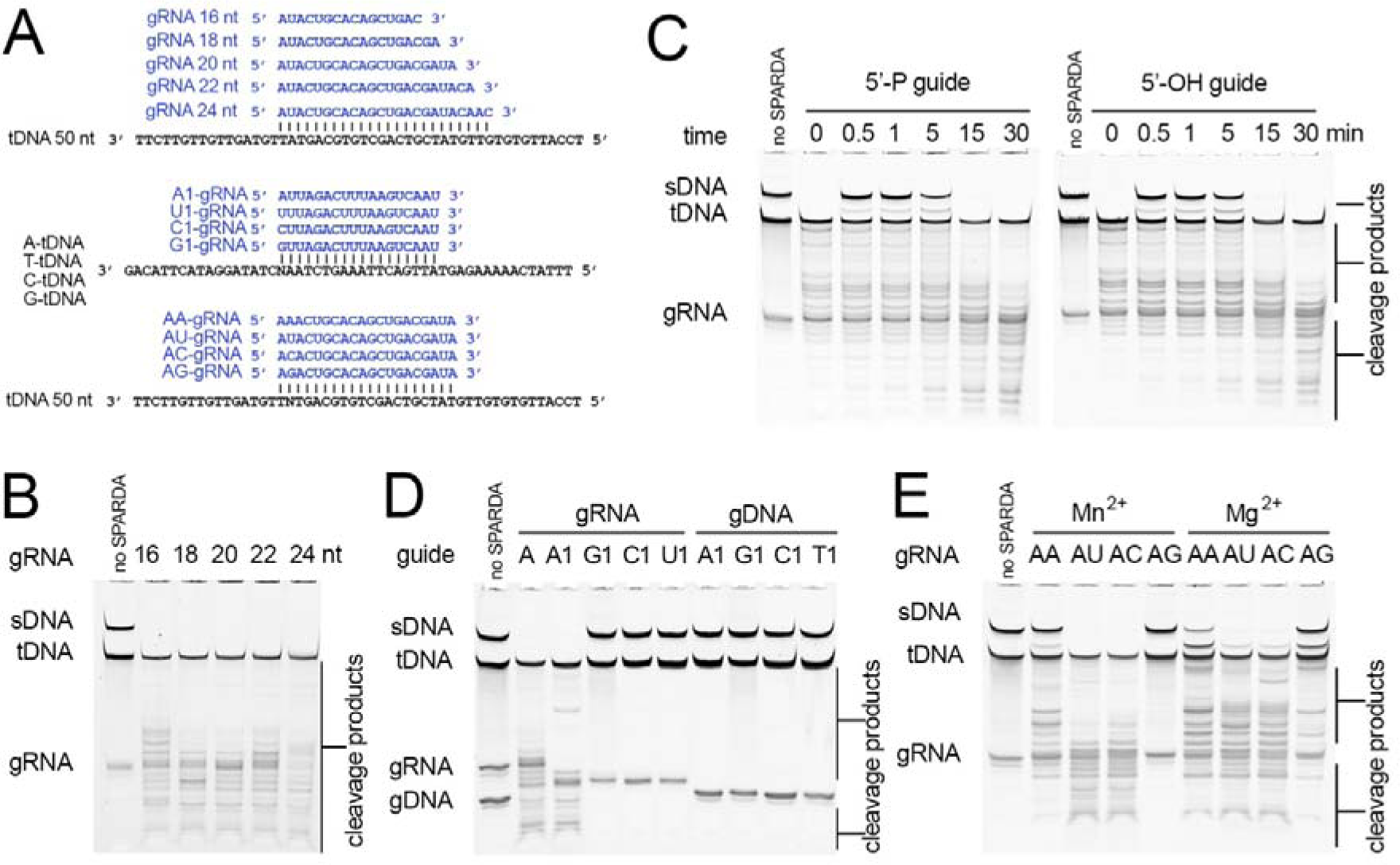
Effects of the gRNA structure on the activity of SPARDA. (A) Structure of gRNAs and single-stranded tDNAs used in the assays. For gRNAs with substitutions of the first or second nucleotides, tDNA contained complementary substitutions at corresponding positions (indicated with ‘N’). (B) Activity of SPARDA with gRNAs of different lengths. (C) Kinetics of substrate DNA cleavage with 5’-phosphorylated and 5’-OH gRNAs. (D) Effects of substitutions of the 5’-nucleotide in gRNA (lanes 2-6) or in gDNA (lanes 7-10) on the activity of SPARDA. SPARDA is activated only by gRNAs with 5’-A (lanes 2 and 3). (E) Effects of substitutions of the second nucleotide in gRNA on the activity of SPARDA. The 5’-end dinucleotides in each gRNA are shown above the gel. All reactions were performed at 30 °C in reaction buffers containing 5 mM Mn^2+^ and 100 mM KCl (or 5 Mg^2+^ and 5 mM KCl in lanes 6-9 in panel E). Positions of gRNA, tDNA, single-stranded collateral substrate DNA (sDNA) and the cleavage products are indicated.

**Extended Data Figure 5.**
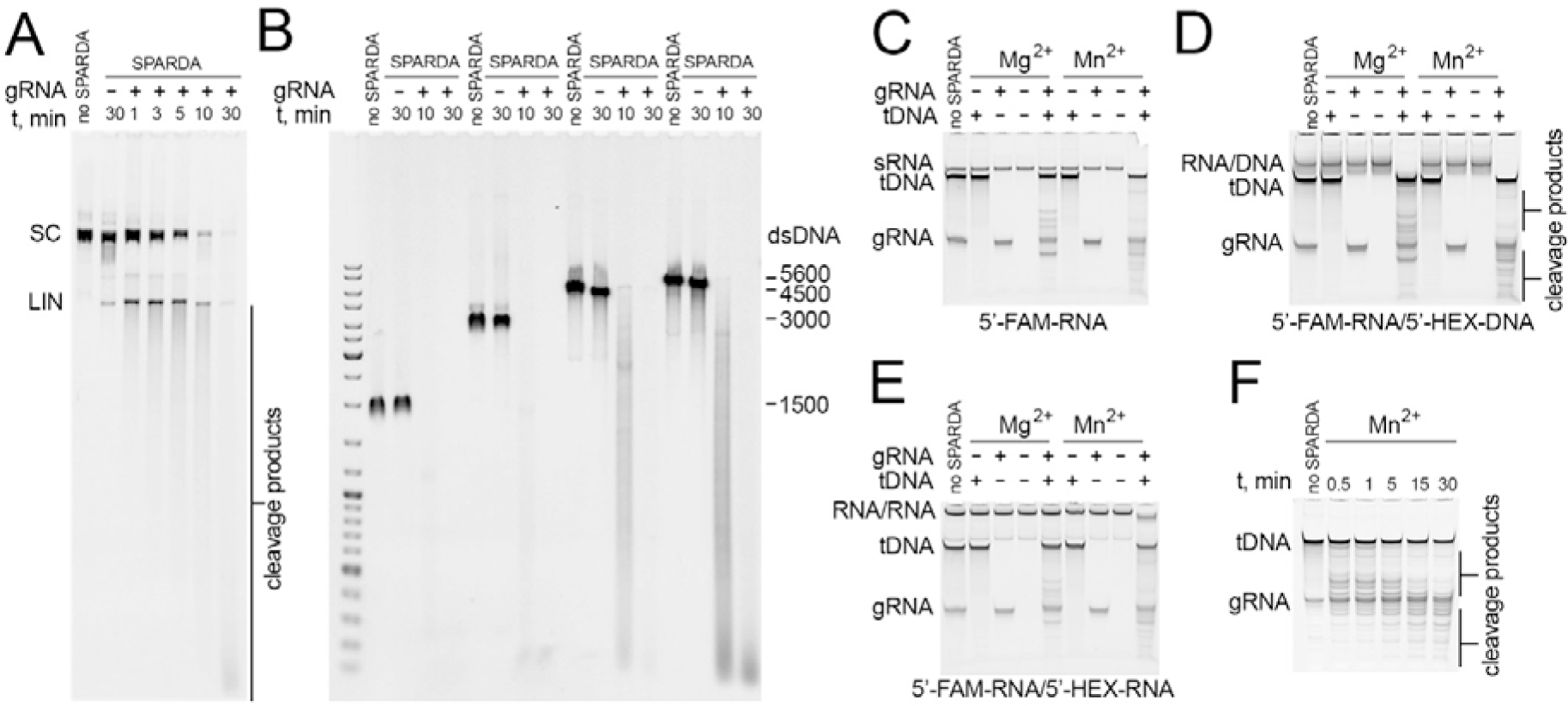
Activity of SPARDA with various collateral substrates. (A) Kinetics of cleavage of supercoiled plasmid DNA (pBAD) by SPARDA. Control samples reactions were performed for 40 minutes without SPARDA (lane 1) or without gRNA (lane 2). (В) Cleavage of linear dsDNA fragments of indicated lengths (obtained after treatment of pBAD by restriction endonucleases with gel purification). The reactions in panels A and B were performed in buffer containing 5 mM Mg^2+^ and 5 mM KCl. (C-E) Analysis of collateral cleavage of 55 nt ssRNA (C), RNA/DNA duplex (D) and dsRNA (E). The reactions were performed with 5’-FAM and 5’-HEX labeled substrate RNA and DNA oligonucleotides, as indicated on the figure, and visualized by SYBR Gold staining. See Table S1 for oligonucleotide sequences. (F) Analysis of gRNA and tDNA cleavage by SPARDA in the absence of collateral substrates. All reactions were performed at 30 °C with wild-type SPARDA (200 nM in panels A and B, 500 nM in panels C-F), with standard 20 nt gRNA and 50 nt single-stranded tDNA in reaction buffers containing 5 mM Mg^2+^ and 5 mM KCl, or 5 mM Mn^2+^ and 100 mM KCl. Positions of gRNA, tDNA, collateral substrates and the cleavage products are indicated.

**Extended Data Figure 6.**
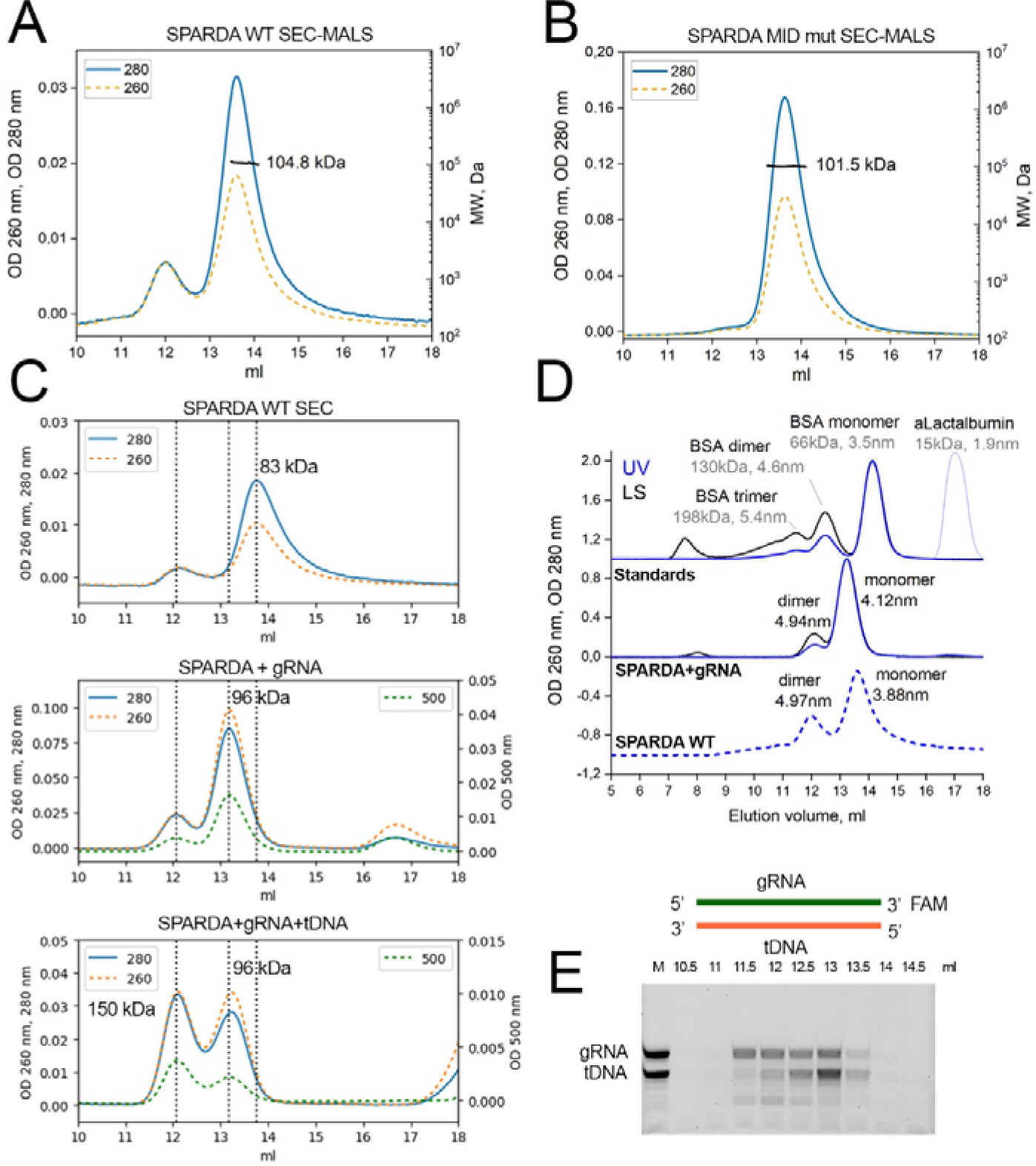
Analysis of the oligomeric state of SPARDA by size-exclusion chromatography and multi-angle light scattering. (A) Analysis of the WT SPARDA complex by SEC-MALS. The absolute molar weight of the NbaAgo/DREN-APAZ heterodimer is indicated. The Mw/Mn polydispersity index was 1.006, suggesting particles with the same Mw across the peak. (B) Analysis of the MID mutant of SPARDA by SEC-MALS, with the absolute molecular weight indicated. The Mw/Mn polydispersity index was 1.000. (C) Size-exclusion chromatography of the purified WT SPARDA complex in the absence of nucleic acids (top), in the presence of 20 nt gRNA (middle) and in the presence of 20 nt gRNA and 20 nt tDNA (bottom). For each sample, OD_280_ and OD_260_ absorbance profiles are shown. OD_500_ profiles are also shown for samples containing gRNA (which was labeled with a FAM residue at its 3’-end, see Table S1). In each panel, positions of monomeric (apparent Mw of 83 or 96 kDa) and dimeric (apparent Mw of ∼150 kDa) SPARDA peaks are indicated. Note that gRNA binding slightly increases the apparent size of SPARDA, as suggested by an increase of the apparent Mw from 83 to 96 kDa and hydrodynamic radius from 3.88 to 4.12 nm, but does not involve any distinct oligomeric state transition. (D) Analysis of the apparent hydrodynamic radii and masses of monomeric (heterodimers of NbaAgo and DREN-APAZ) and dimeric (dimers of heterodimers) SPARDA complexes, based on calibration with standard albumin samples. (E) Analysis of nucleic acids from various fractions after size-exclusion chromatography of the SPARDA sample pre-incubated with gRNA and tDNA (from the bottom profile in panel C). Based on the OD_260_/OD_280_ ratio and the electrophoretic analysis of gRNA and tDNA in each fraction it can be concluded that dimers of the heterodimeric SPARDA complex contain bound gRNA, while monomeric heterodimers contain both gRNA and tDNA. The dimeric peak of SPARDA is observed in the presence of gRNA (middle panel), gRNA and tDNA (bottom panel). The presence of a minor dimeric fraction in WT SPARDA (panel C, top, and panel D, bottom) likely results from binding of gRNAs co-purified with SPARDA (see Extended data Fig. 10A), in contrast to the gRNA-free main monomeric peak.

**Extended Data Figure 7.**
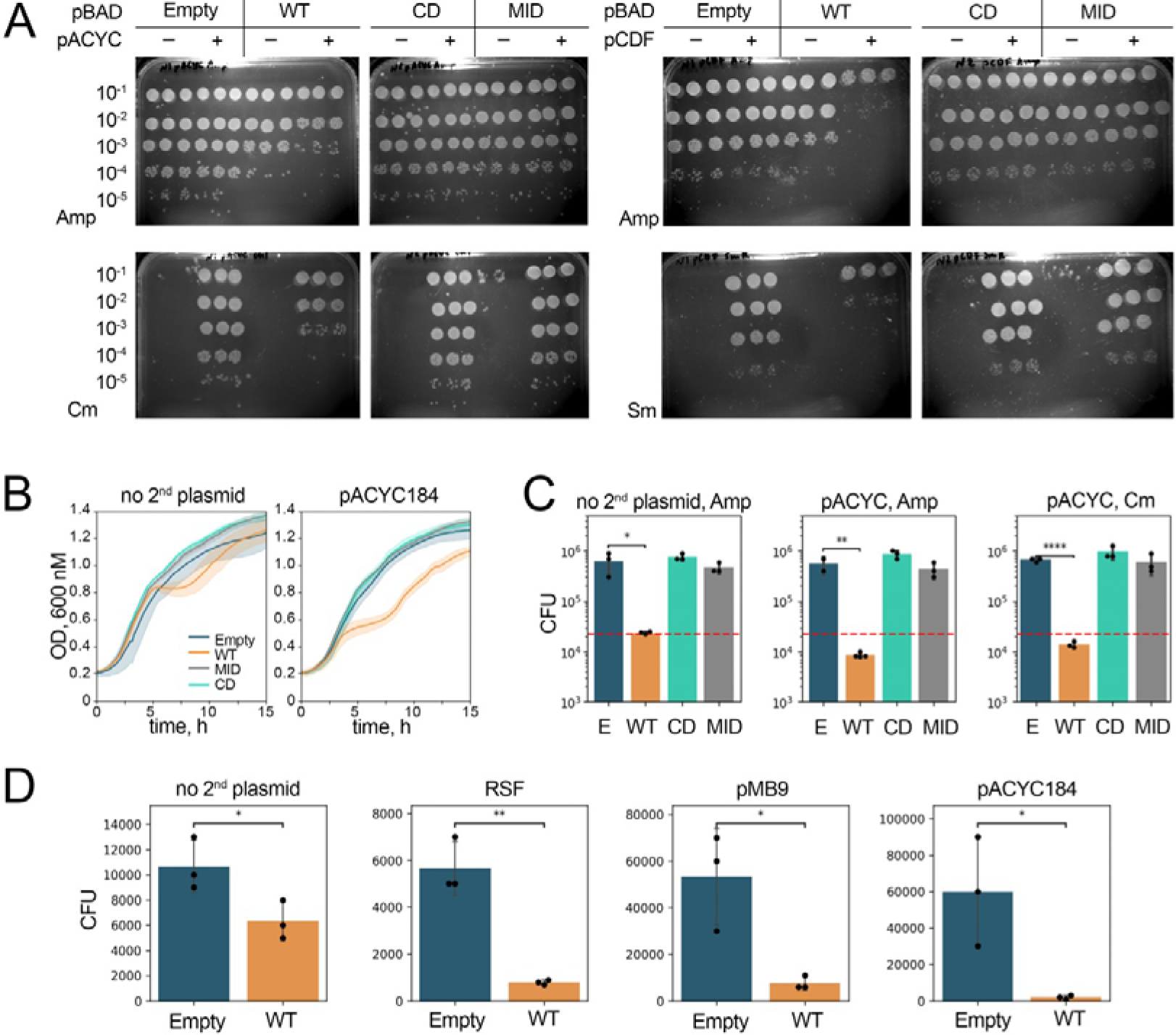
Plasmid interference induced by SPARDA. (A) Measurements of CFU numbers for *E. coli* strains containing empty pBAD (Amp^R^) or pBAD encoding SPARDA (WT or mutant, CD or MID), and transformed with pACYC (Cm^R^, left) or pCDF (Sm^R^, right) interfering plasmids. The cells were grown for 2.5 hours at 37 °C in the presence of Amp and 0.1% Ara, to induce SPARDA, serial dilutions were plated on LB agar containing Amp or Cm or Sm and 1% Glc (to repress SPARDA expression), and CFU numbers were counted after overnight growth. The calculation of the CFU numbers is shown in Fig. 3C. (B) Growth of *E. coli* strains expressing wild-type (WT) or mutant SPARDA (MID, substitutions in the MID domain; CD, catalytically dead) from pBAD, or containing control empty pBAD, depending on the presence of the second interfering plasmid (pACYC184). Means and standard deviations from 6 replicate experiments. (C) Numbers of viable cells expressing SPARDA (WT or its MID and CD mutants) or containing empty pBAD (‘E’), depending on the presence of the pACYC184 plasmid. Means and standard deviations from 3 replicate experiments. (D) Numbers of viable cells containing the WT SPARDA plasmid (or the control empty pBAD) and additional interfering plasmids (RSF1010, pMB9, pACYC184, or no second plasmid). The cells were grown in the presence of 0.1% Ara to induce SPARDA expression. Statistically significant changes in the CFU numbers are indicated (*p<0.05; **p<0.01, ****p<0.0001).

**Extended Data Figure 8.**
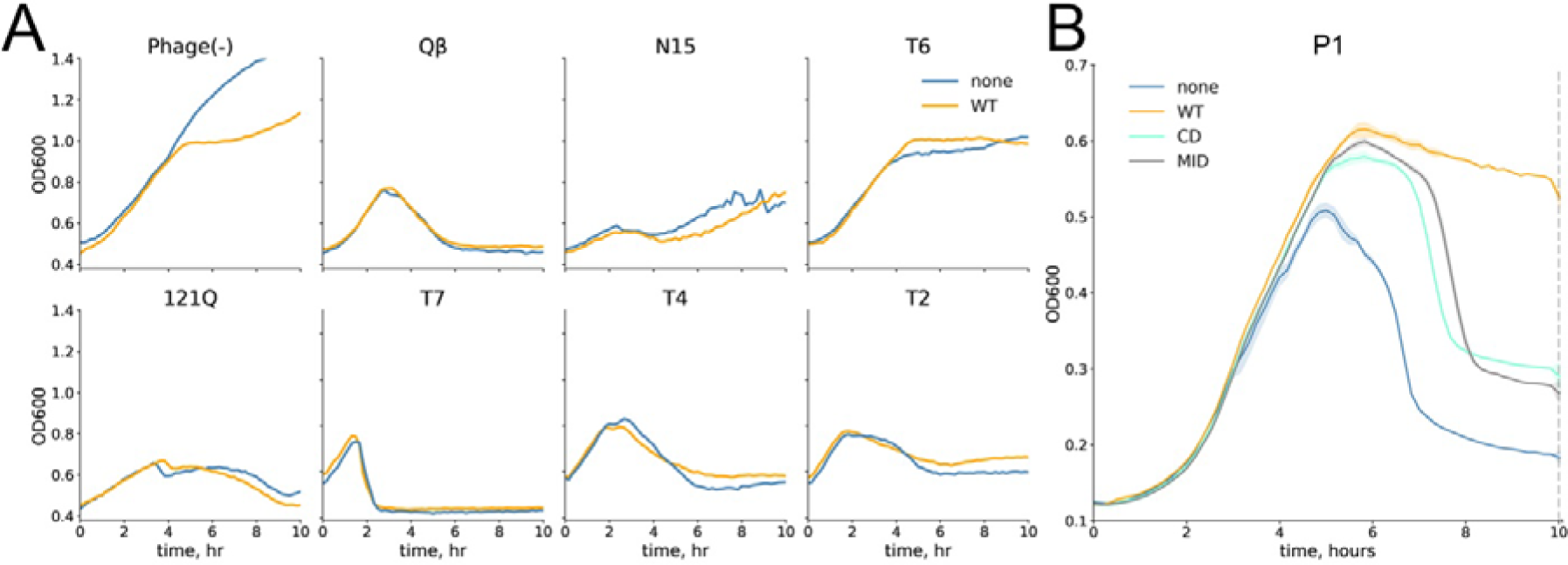
Effects of SPARDA expression on phage infection. (A) Growth and collapse of *E. coli* cultures (MG1655) infected with various phages, expressing WT SPARDA or containing the control empty pBAD. Single representative experiments for each phage are shown. (B) Growth of *E. coli* BL21(DE3) infected with phage P1, expressing WT or mutant SPARDA variants. Averages from three independent experiments performed in parallel are shown. These cultures were used for determination of the PFU and CFU numbers (for the 9 hour time point indicated with a dashed line) (Fig. 3F). For sequencing of small and long RNAs (Fig. 3I), the cultures were grown under similar conditions and the samples were taken at 5 hours, just before phage-induced lysis. All strains were grown in the presence of 0.1% Ara to induce SPARDA expression.

**Extended Data Figure 9.**
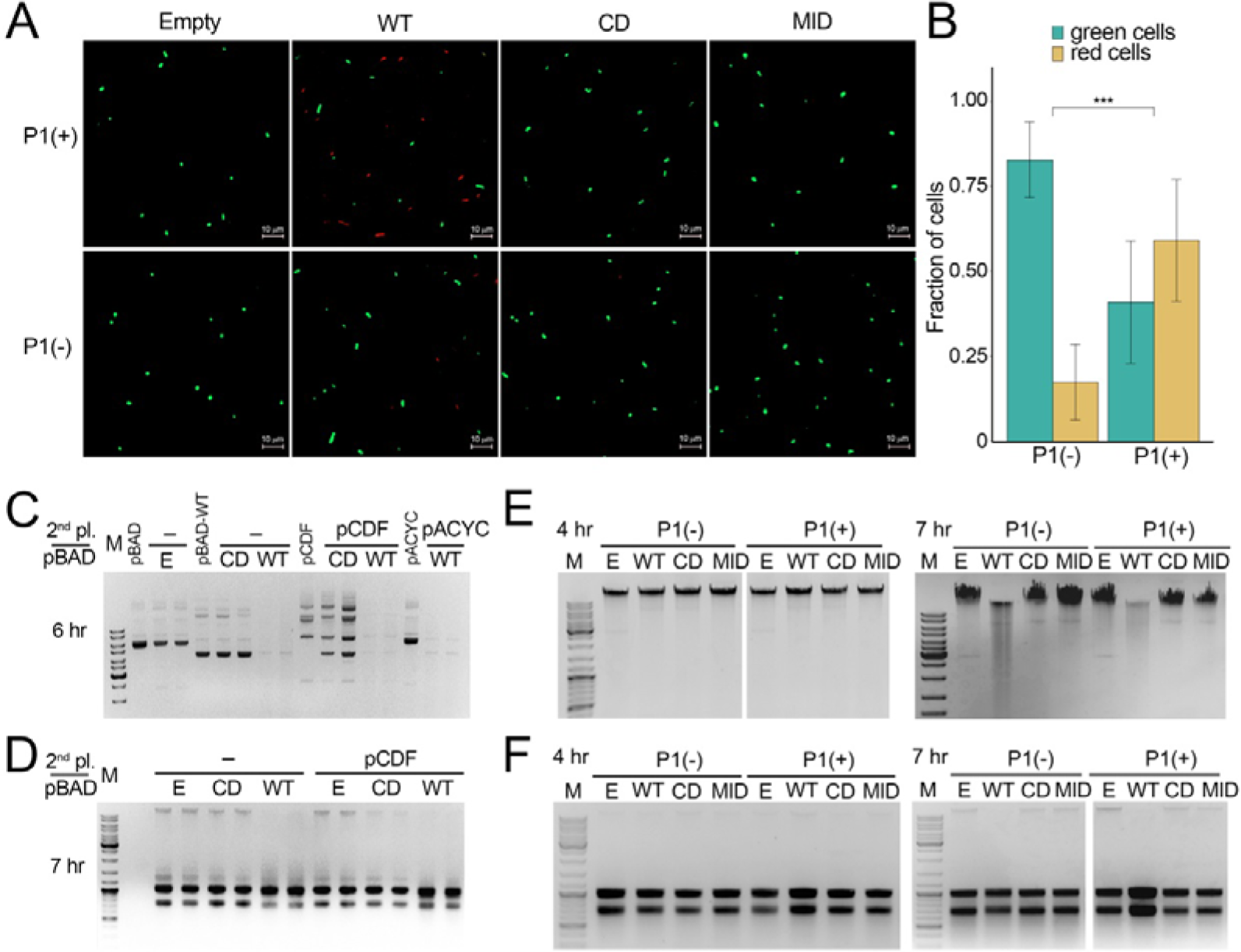
Analysis of the effects of SPARDA on cell viability and degradation of DNA and RNA *in vivo*. (A) Analysis of cell viability by propidium iodide (PI) staining. Cell cultures expressing wild-type or mutant SPARDA or containing the control empty pBAD were infected with phage P1 (MOI 3.6×10^-4^), grown for 14 hours, stained with PI and Syto 9 and analyzed by microscopy. Representative fields of view are shown. The scale bar is 10 μm. (B) Quantification of the proportion of live (green) and dead (red) cells in *E. coli* cultures expressing SPARDA in the absence and in the presence of phage P1. (C) Analysis of plasmid DNA integrity. Plasmid DNA was purified from *E. coli* strains containing pBAD (empty ‘E’, or encoding SPARDA, WT or CD), or pBAD together with pACYC or pCDF, at 6 h of growth after induction of SPARDA (0.1% Ara). The results of two replicate experiments are shown. (D) Analysis of total RNA purified from *E. coli* cultures containing pBAD (empty ‘E’, or encoding WT or CD SPARDA), or pBAD together with pCDF, at 7 h of growth after induction of SPARDA. (E) Analysis of genomic DNA integrity depending on phage infection. Chromosomal DNA was purified from *E. coli* cultures expressing WT or mutant SPARDA from pBAD plasmids, or containing the control empty pBAD, in the absence and in the presence of phage P1, at 4 and 7 h after induction of SPARDA and P1 infection. (F) Analysis of total RNA purified from *E. coli* during phage infection. Total RNA was isolated from the same strains as in panel E, at 4 and 7 hours of cell growth in the absence of phages and during infection with phage P1.

**Extended Data Figure 10.**
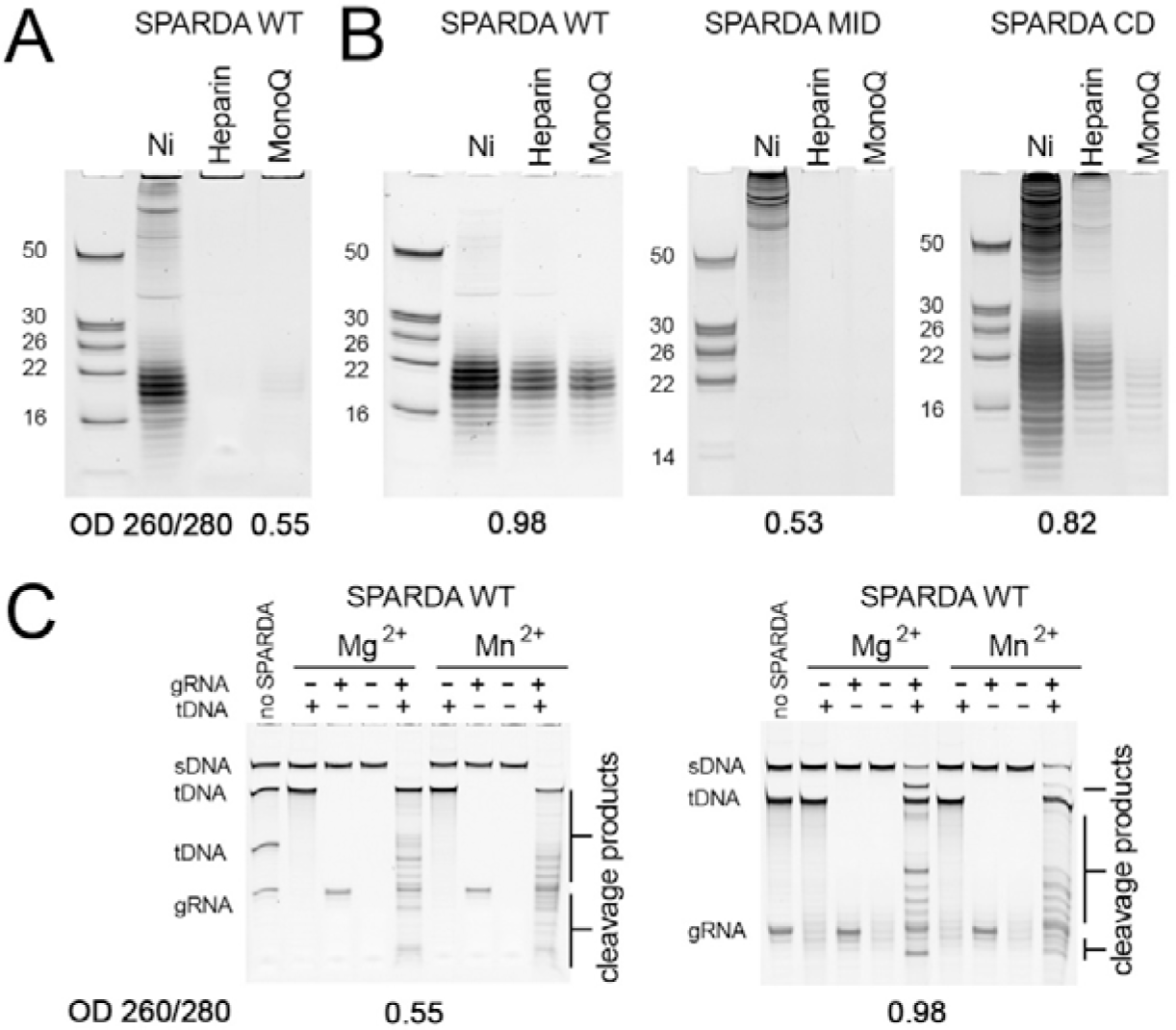
Purification of small guide RNAs associated with NbaSPARDA in *E. coli*. (A) Small RNAs associated with WT SPARDA at different steps of the standard purification protocol (Ni-chelating, heparin and MonoQ columns). Most gRNAs are removed after heparin chromatography. The OD_260/280_ ratio of the final sample is 0.55. (B) Small RNAs associated with WT and mutant SPARDA variants at different steps of purification using a modified protocol, in which EDTA was omitted from the buffers during heparin and anion-exchange chromatography. Significant amounts of gRNAs remain associated within the WT and CD complexes (but not the MID mutant) through all purification steps, indicative of stable gRNA binding. The OD_260/280_ ratios of the final WT, MID and CD samples are 0.98, 0.53 and 0.82, respectively. (C) Comparison of the activities of WT SPARDA purified by different protocols. Preparations of SPARDA containing cell-derived gRNAs (OD_260/280_=0.98, top) has a lower nuclease activity in comparison with guide-free SPARDA (OD_260/280_=0.55, bottom), when activated by specific gRNA (20 nt) and single-stranded tDNA (50 nt).

**Extended Data Table 1.**
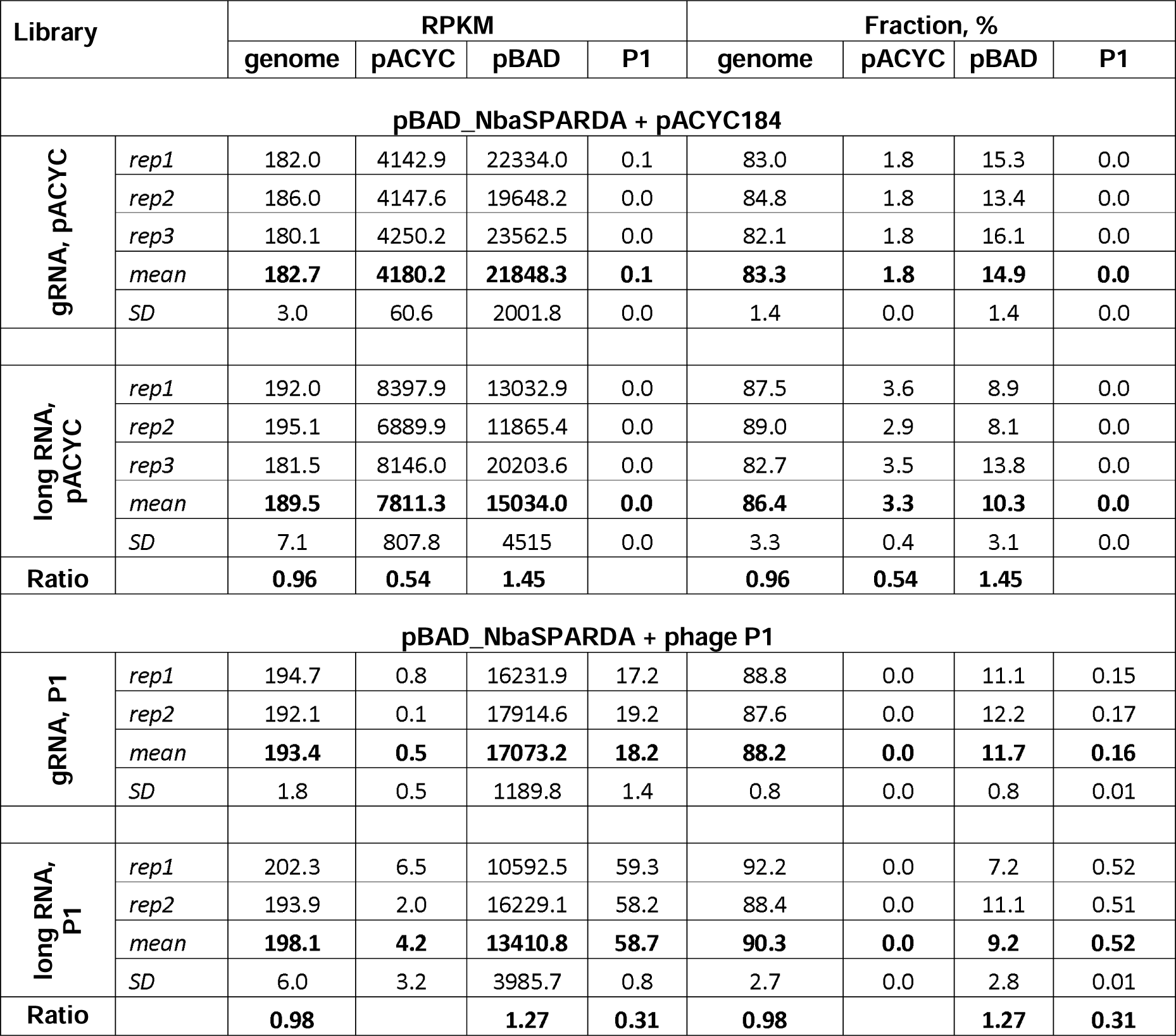
Distribution of gRNA and long RNA reads between chromosomal, plasmid and phage sequences in *E. coli* strains expressing NbaSPARDA.

| Library |  | RPKM |  |  |  | Fraction, % |  |  |  |
| --- | --- | --- | --- | --- | --- | --- | --- | --- | --- |
|  |  | genome | pACYC | pBAD | P1 | genome | pACYC | pBAD | P1 |
| **pBAD_NbaSPARDA + pACYC184** |  |  |  |  |  |  |  |  |  |
| gRNA, pACYC | *rep1* | 182.0 | 4142.9 | 22334.0 | 0.1 | 83.0 | 1.8 | 15.3 | 0.0 |
|  | *rep2* | 186.0 | 4147.6 | 19648.2 | 0.0 | 84.8 | 1.8 | 13.4 | 0.0 |
|  | *rep3* | 180.1 | 4250.2 | 23562.5 | 0.0 | 82.1 | 1.8 | 16.1 | 0.0 |
|  | *mean* | **182.7** | **4180.2** | **21848.3** | **0.1** | **83.3** | **1.8** | **14.9** | **0.0** |
|  | *SD* | 3.0 | 60.6 | 2001.8 | 0.0 | 1.4 | 0.0 | 1.4 | 0.0 |
| long RNA, pACYC | *rep1* | 192.0 | 8397.9 | 13032.9 | 0.0 | 87.5 | 3.6 | 8.9 | 0.0 |
|  | *rep2* | 195.1 | 6889.9 | 11865.4 | 0.0 | 89.0 | 2.9 | 8.1 | 0.0 |
|  | *rep3* | 181.5 | 8146.0 | 20203.6 | 0.0 | 82.7 | 3.5 | 13.8 | 0.0 |
|  | *mean* | **189.5** | **7811.3** | **15034.0** | **0.0** | **86.4** | **3.3** | **10.3** | **0.0** |
|  | *SD* | 7.1 | 807.8 | 4515 | 0.0 | 3.3 | 0.4 | 3.1 | 0.0 |
| Ratio |  | **0.96** | **0.54** | **1.45** |  | **0.96** | **0.54** | **1.45** |  |
| **pBAD_NbaSPARDA + phage P1** |  |  |  |  |  |  |  |  |  |
| gRNA, P1 | *rep1* | 194.7 | 0.8 | 16231.9 | 17.2 | 88.8 | 0.0 | 11.1 | 0.15 |
|  | *rep2* | 192.1 | 0.1 | 17914.6 | 19.2 | 87.6 | 0.0 | 12.2 | 0.17 |
|  | *mean* | **193.4** | **0.5** | **17073.2** | **18.2** | **88.2** | **0.0** | **11.7** | **0.16** |
|  | *SD* | 1.8 | 0.5 | 1189.8 | 1.4 | 0.8 | 0.0 | 0.8 | 0.01 |
| long RNA, P1 | *rep1* | 202.3 | 6.5 | 10592.5 | 59.3 | 92.2 | 0.0 | 7.2 | 0.52 |
|  | *rep2* | 193.9 | 2.0 | 16229.1 | 58.2 | 88.4 | 0.0 | 11.1 | 0.51 |
|  | *mean* | **198.1** | **4.2** | **13410.8** | **58.7** | **90.3** | **0.0** | **9.2** | **0.52** |
|  | *SD* | 6.0 | 3.2 | 3985.7 | 0.8 | 2.7 | 0.0 | 2.8 | 0.01 |
| Ratio |  | **0.98** |  | **1.27** | **0.31** | **0.98** |  | **1.27** | **0.31** |

**Extended Data Figure 11.**
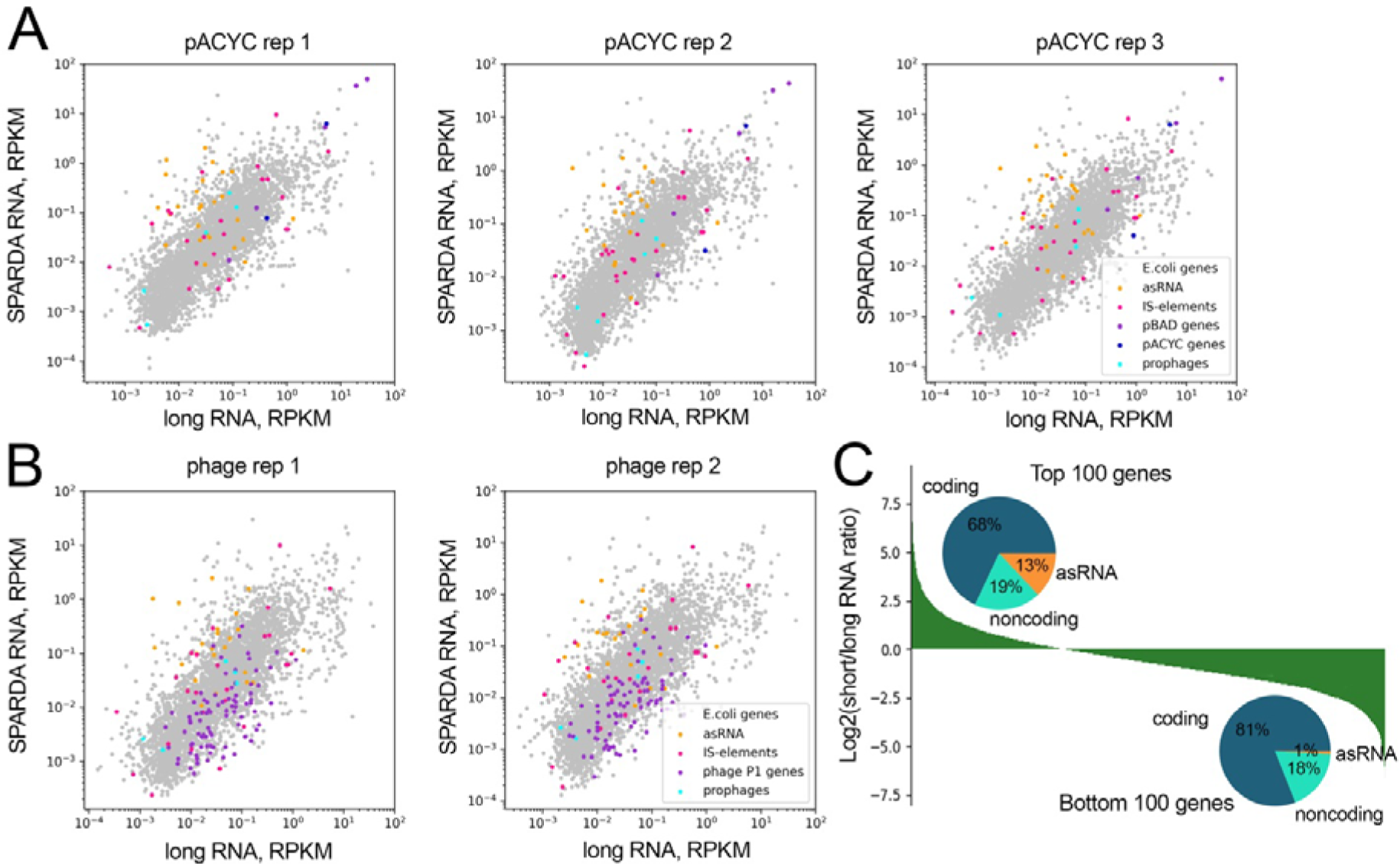
(A) Correlation between the abundance of gRNAs and the RNA transcriptome of *E. coli* strains expressing WT SPARDA from pBAD and containing pACYC (*top*) or infected with phage P1 (*bottom*). Individual replicate experiments for Fig. 3I are presented. Plasmid genes (pBAD and pACYC) are shown in blue and violet; phage P1 genes are shown in magenta. Regulatory antisense RNAs, prophage genes, and IS elements are shown in orange, pink and turquoise, respectively. (B) Distribution of various types of transcripts among the genes which are enriched or depleted for gRNAs. For all *E. coli* genes, the ratios of small gRNAs to long RNA transcripts were calculated, and the genes were sorted by this ratio in descending order (from left to right). The distributions of coding, noncoding and regulatory antisense transcripts among the top and bottom 100 genes are shown in circular diagrams.

**Extended Data Figure 12.**
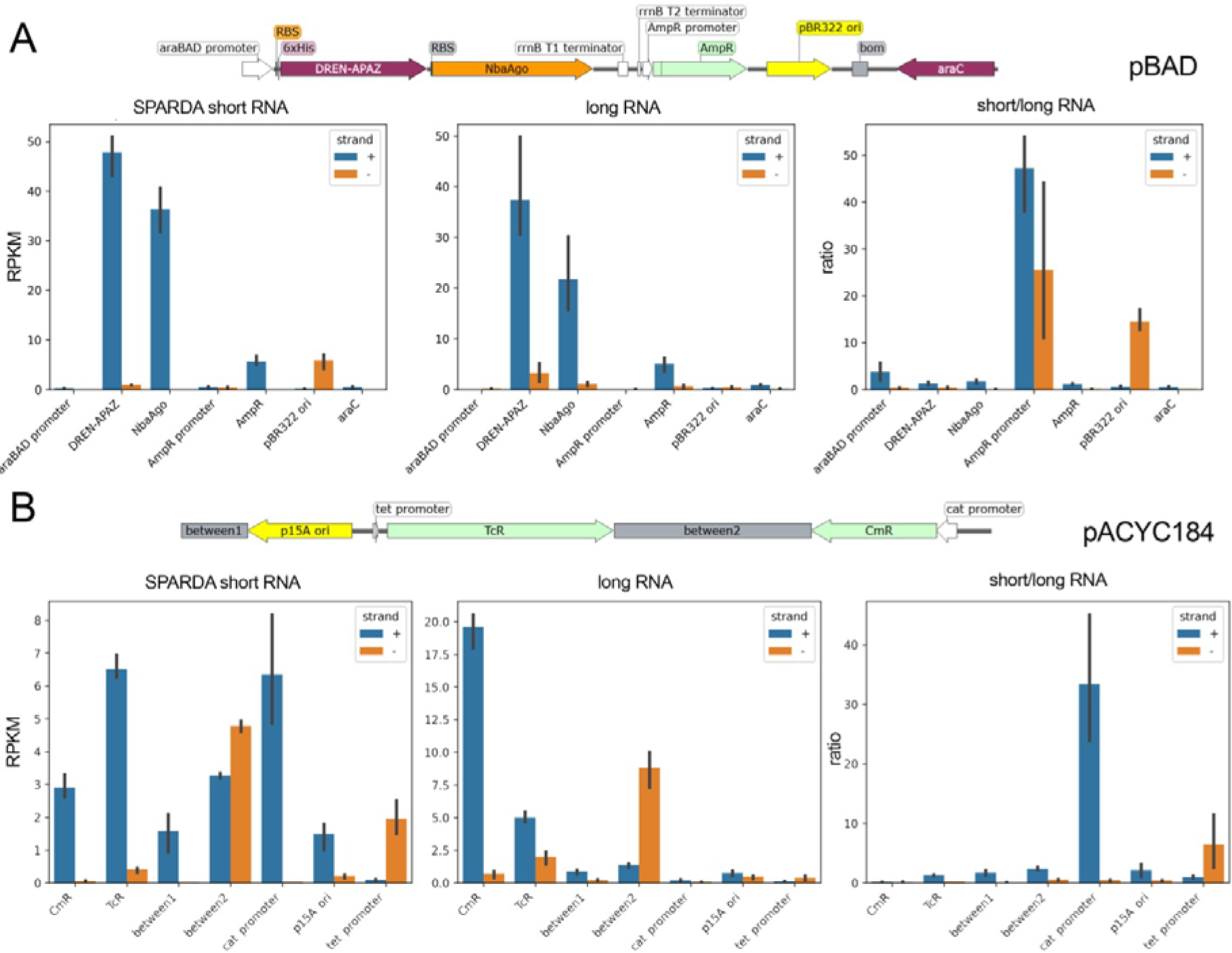
Analysis of SPARDA gRNAs and long RNAs for plasmid DNA. (A) pBAD plasmid. (B) pACYC184 plasmid. SPARDA-associated short gRNAs and total RNA were isolated from *E. coli* containing pBAD_NbaSPARDA and pACYC184 at 5 h after induction of SPARDA, sequenced and mapped to the plasmids. The amounts of short gRNAs and long RNA transcripts are shown in RPKM (reads per kilobase per million of aligned reads in the library) for each DNA strand and plotted for each plasmid region (*left* and *middle* panels). The ratio of short to long RNAs is shown on the *right* panels. Means and standard deviations from three replicate experiments.

**Extended Data Figure 13.**
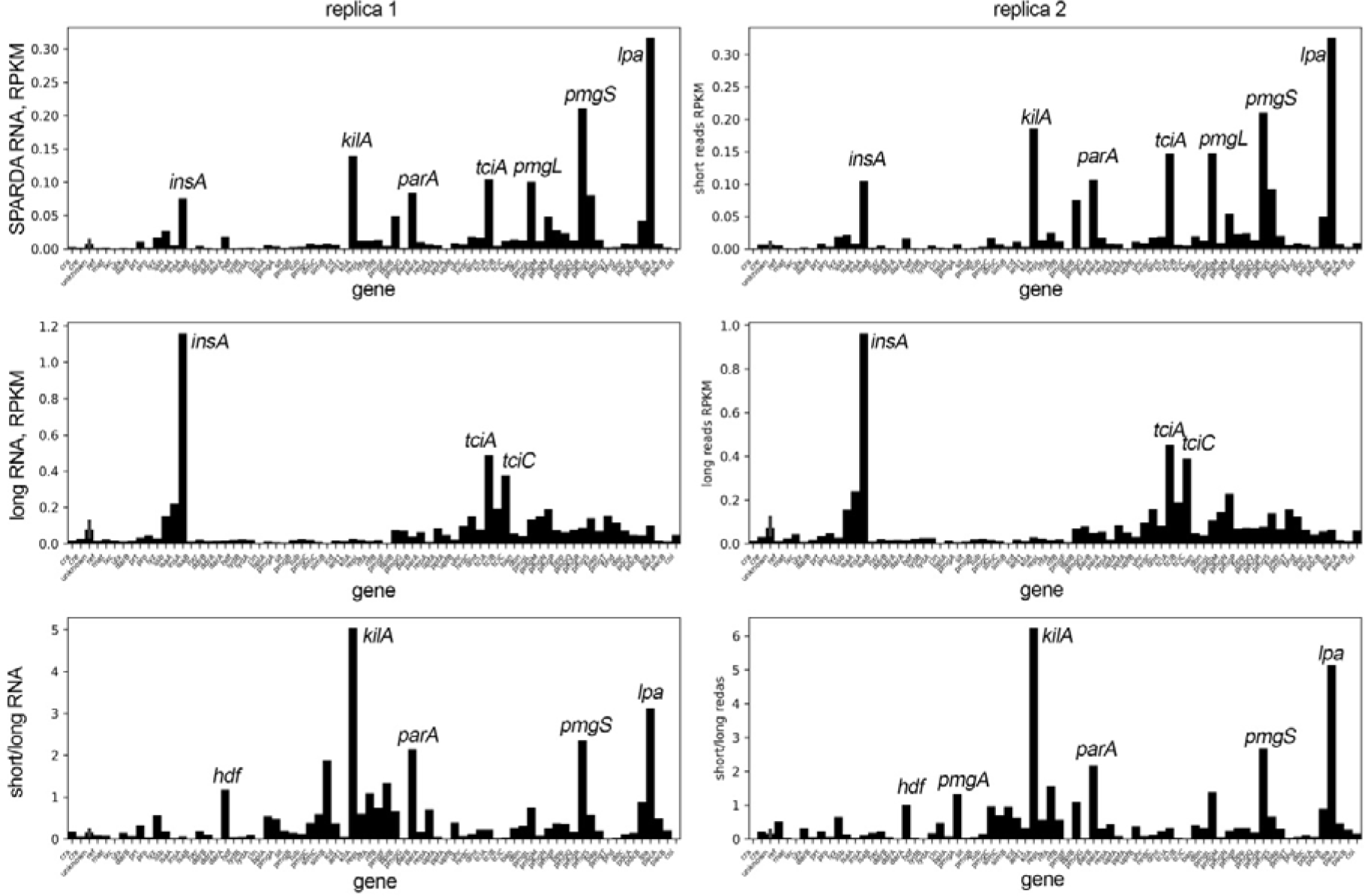
Analysis of SPARDA gRNAs and long RNAs for the phage P1 genome. SPARDA-associated short gRNAs and total RNA were isolated from *E. coli* at 5 hours of infection with phage P1, sequenced and mapped to the phage genome (GenBank ID GCA_025787755.1). The amounts of short gRNAs and long RNA transcripts are shown in RPKM for each phage gene (*top* and *middle* panels, respectively). The ratio of short to long RNAs is shown on the *bottom* panels. The data from two replicate experiments. Genes with the highest amounts of short gRNAs, long RNA transcripts and the highest ratio of short to long RNAs are labeled. Regions enriched in short RNAs in both replicas include *hdf* (part of the phage antirestriction system), *kilA* (interferes with plasmid establishment after transformation or is lethal to cells), *parA* (part of the partitioning system), *pmgS* (putative morphogenetic function) and *lpa* (transcription activator of late genes).

**Extended Data Figure 14.**
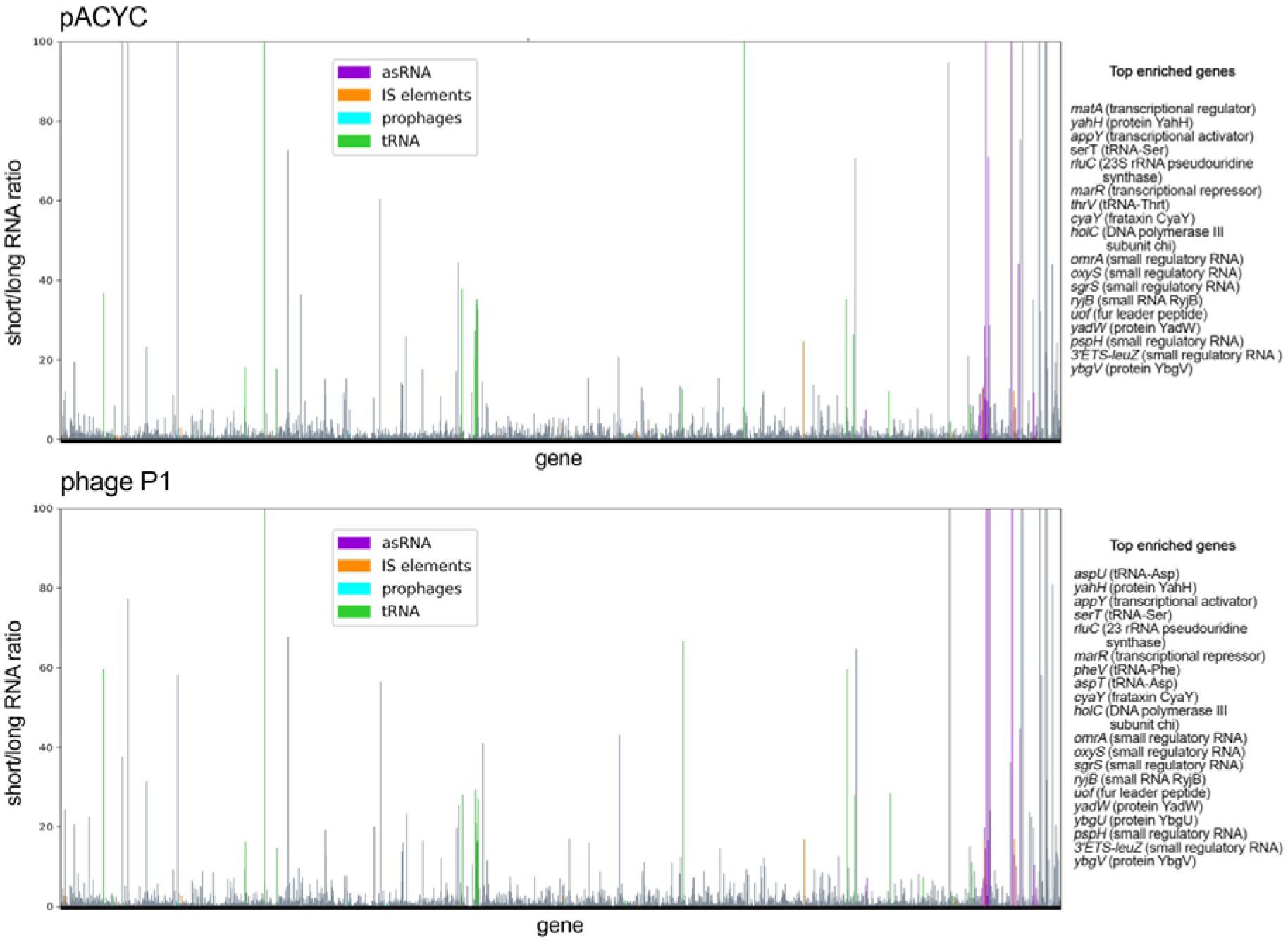
Analysis of SPARDA gRNAs and long RNAs for the *E. coli* chromosome. SPARDA-associated short gRNAs and total RNA were isolated from *E. coli* cells expressing SPARDA from pBAD_NbaSPARDA and containing pACYC184 (*top*) or infected with phage P1 (*bottom*), sequenced and mapped to the MG1655 reference genome (GenBank ID GCA_000005845.2). The amounts of short gRNAs and long RNA transcripts were expressed in RPKM for each *E. coli* gene and their ratio was calculated. The sequences corresponding to rRNA were omitted from analysis. Means from three (for the *E. coli* strain containing pACYC184) and two (for *E. coli* infected with phage P1) replicate experiments are shown on the upper and bottom panels, respectively. The list of top genes with the highest ratio of short to long RNAs is shown for each averaged plot.

## Supplementary Information

**Table S1.**
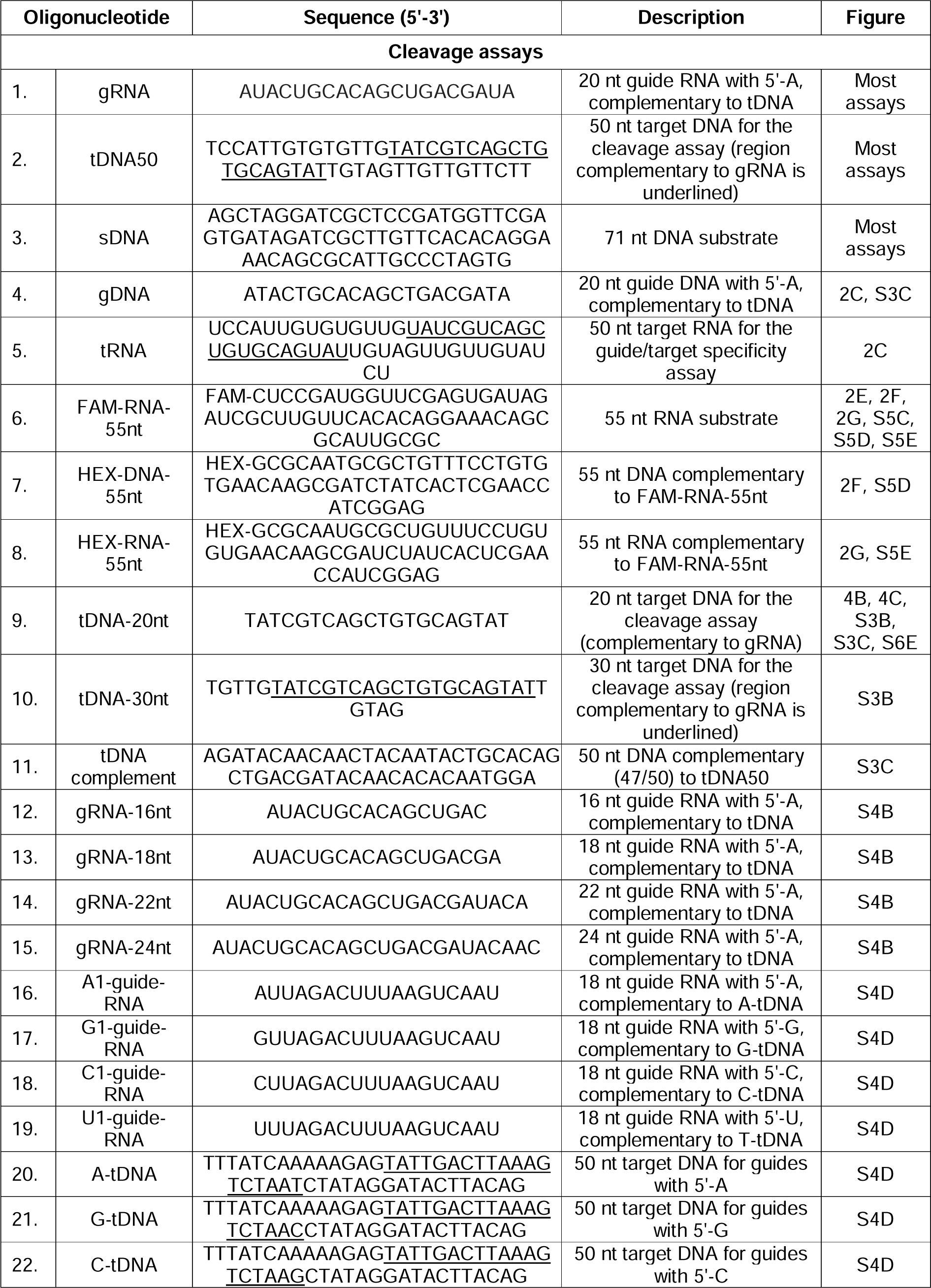
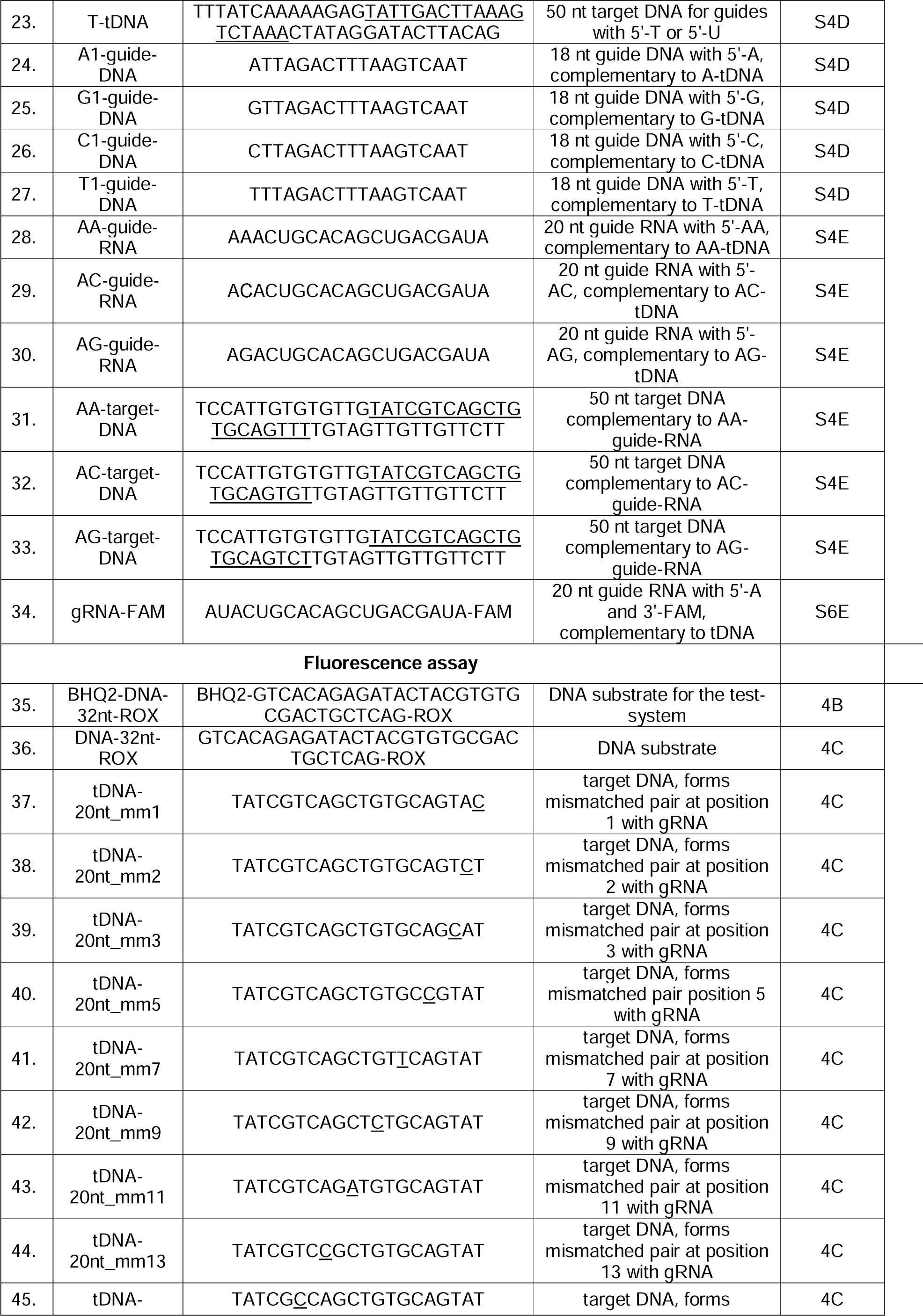
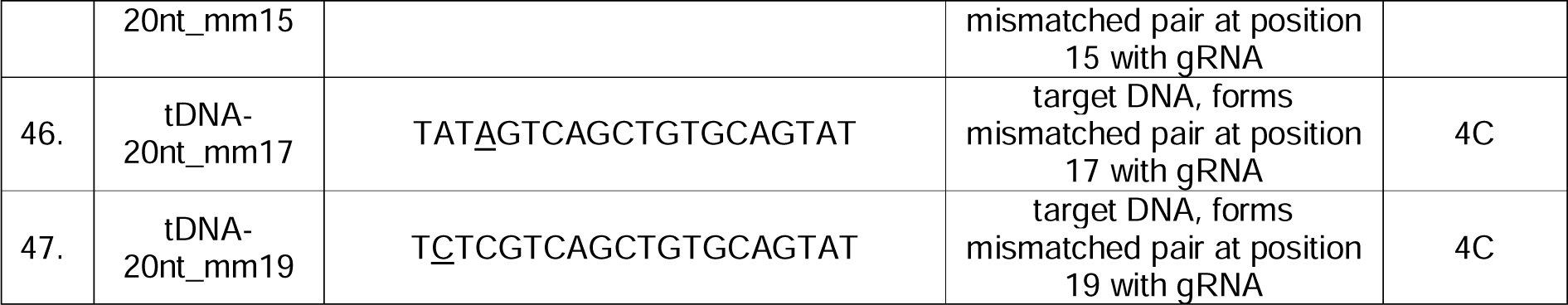
Oligonucleotides used in this study.

**Table S2.** Small RNA and total RNA libraries. The libraries were obtained from *E. coli* strains expressing SPARDA from pBAD and co-transformed with pACYC184 or infected with phage P1.

| Library type | Strain | Plasmid/phage | Growth Conditions | Growth Phase | Library type | BioProject accession number/BioSample accession number | SRA accession number |
| --- | --- | --- | --- | --- | --- | --- | --- |
| **Small RNAs** | *E. coli* BL21(DE3) | pACYC184 | LB +MgCl<sub>2</sub>+CaCl<sub>2</sub>+Ampicillin, arabinose induction, 30°C | 5h after induction | smRNA, SE50 | PRJNA975910/<br>SAMN35348438 | TBA |
|  | *E. coli* BL21(DE3) | pACYC184 | LB +MgCl<sub>2</sub>+CaCl<sub>2</sub>+Ampicillin, arabinose induction, 30°C | 5h after induction | smRNA, SE50 | PRJNA975910/<br>SAMN35348439 | TBA |
|  | *E. coli* BL21(DE3) | pACYC184 | LB +MgCl<sub>2</sub>+CaCl<sub>2</sub>+Ampicillin, arabinose induction, 30°C | 5h after induction | smRNA, SE50 | PRJNA975910/<br>SAMN35348440 | TBA |
|  | *E. coli* BL21(DE3) | Phage P1 | LB +MgCl<sub>2</sub>+CaCl<sub>2</sub>+Ampicillin, arabinose induction, 30°C | 5h after induction and infection | smRNA, SE50 | PRJNA975910/<br>SAMN35348441 | TBA |
|  | *E. coli* BL21(DE3) | Phage P1 | LB +MgCl<sub>2</sub>+CaCl<sub>2</sub>+Ampicillin, arabinose induction, 30°C | 5h after induction and infection | smRNA, SE50 | PRJNA975910/<br>SAMN35348442 | TBA |
| **Total RNA** | *E. coli* BL21(DE3) | pACYC184 | LB +MgCl<sub>2</sub>+CaCl<sub>2</sub>+Ampicillin, arabinose induction, 30°C | 5h after induction | tRNA, SE50 | PRJNA975910/<br>SAMN35348438 | TBA |
|  | *E. coli* BL21(DE3) | pACYC184 | LB +MgCl<sub>2</sub>+CaCl<sub>2</sub>+Ampicillin, arabinose induction, 30°C | 5h after induction | tRNA, SE50 | PRJNA975910/<br>SAMN35348439 | TBA |
|  | *E. coli* BL21(DE3) | pACYC184 | LB +MgCl<sub>2</sub>+CaCl<sub>2</sub>+Ampicillin, arabinose induction, 30°C | 5h after induction | tRNA, SE50 | PRJNA975910/<br>SAMN35348440 | TBA |
|  | *E. coli* BL21(DE3) | Phage P1 | LB +MgCl<sub>2</sub>+CaCl<sub>2</sub>+Ampicillin, arabinose induction, 30°C | 5h after induction and infection | tRNA, SE50 | PRJNA975910/<br>SAMN35348441 | TBA |
|  | *E. coli* BL21(DE3) | Phage P1 | LB +MgCl<sub>2</sub>+CaCl<sub>2</sub>+Ampicillin, arabinose induction, 30°C | 5h after induction and infection | tRNA, SE50 | PRJNA975910/<br>SAMN35348442 | TBA |

